# SMG6-dependent RNA decay maintains dsRNA homeostasis and restrains immunogenicity in hepatocellular carcinoma

**DOI:** 10.1101/2025.01.13.632668

**Authors:** Enes S. Arpa, Virginie Ricci, Ilja E. Shapiro, Marija Jokic, Yahya Mohammadzadeh, Lisa Bertrand, Irene Gonzalez-Menendez, Leticia Quintanilla-Martinez, Alexandre P. Bénéchet, Dirk Mossmann, Michael N. Hall, Jan Rehwinkel, Giovanni Ciriello, Tatiana V. Petrova, Sanjiv A. Luther, Michal Bassani-Sternberg, David Gatfield

## Abstract

Nonsense-mediated mRNA decay (NMD) eliminates aberrant transcripts to maintain transcriptome integrity, yet its broader physiological roles remain incompletely defined. Here, using a liver-specific, inducible mouse model, we selectively inactivate the endonuclease SMG6 – the terminal effector of NMD – in a genetic model of hepatocellular carcinoma (HCC). SMG6 loss completely prevents tumour development and triggers robust innate and adaptive immune responses. Mechanistically, SMG6 inactivation causes the cytoplasmic accumulation of endogenous double-stranded RNAs (dsRNAs), leading to type I interferon induction via the dsRNA sensor MDA5 and revealing an unexpected physiological role for NMD in maintaining dsRNA homeostasis. In parallel, stabilisation and translation of normally degraded transcripts generate non-canonical MHC-I-presented peptides that elicit potent CD8^+^ T-cell responses. These findings identify SMG6-dependent NMD as a central gatekeeper that restrains dsRNA-driven antiviral signalling and suppresses immunogenic transcript expression. Our work establishes selective SMG6 inhibition as a promising strategy to enhance tumour immunogenicity and suggests that targeting terminal NMD activity could represent a conceptually distinct avenue for cancer immunotherapy.

## Introduction

Eukaryotic cells employ nonsense-mediated mRNA decay (NMD) as a critical post-transcriptional quality control mechanism that degrades transcripts harbouring premature termination codons (PTCs), thereby preventing the accumulation of truncated, potentially toxic proteins ^1, 2^. PTCs typically arise from mutations, transcriptional or splicing errors. In addition to its canonical surveillance function, NMD is increasingly recognised as a broader regulator of gene expression, shaping the transcriptome through the selective degradation of physiological mRNAs ^3^. NMD is carried out by a multiprotein machinery centred on the helicase UPF1 and culminates in an endonucleolytic cleavage step executed by the PilT N-terminus (PIN)-domain nuclease SMG6, whose active site is complemented by SMG5 ^4–6^.

In cancer, the role of NMD is multifaceted and context-dependent ^7^. On the one hand, it may suppress tumorigenesis by limiting the expression of mutated, potentially oncogenic transcripts. On the other, it can promote immune evasion by eliminating mRNAs that encode neoantigens, particularly those arising from frameshift-inducing insertion/deletion mutations that are common in cancer genomes. By suppressing such immunogenic RNAs, NMD may dampen anti-tumour immune surveillance and reduce responsiveness to immunotherapies ^8–12^.

Previous studies investigating NMD in cancer have primarily relied on constitutive depletion of core NMD components such as UPF1 or SMG proteins in cell lines or transplantable tumour models ^13–17^. While these approaches have provided important insights, they are limited in their ability to capture early tumorigenic events and the dynamics of immune surveillance *in vivo*. Moreover, several NMD factors – including UPF1, SMG1 and SMG6 – have well-established functions independent of mRNA decay, notably in genome stability, DNA replication and telomere maintenance ^18^. As a result, global loss of these proteins can introduce additional cellular defects that complicate the attribution of observed phenotypes specifically to impaired NMD. Importantly, prior work has shown that the endonuclease activity of SMG6 can be genetically uncoupled from its non-NMD roles ^19^, enabling selective interrogation of its NMD function without disrupting SMG6’s broader cellular activities.

To address these limitations, we developed a liver-specific, inducible mouse model that selectively disables the endonuclease activity of SMG6 without ablating the protein entirely ^20^. We combined this model with a genetic background of hepatocellular carcinoma (HCC) that closely mirrors human disease progression, driven by hepatic deletion of *Pten* and *Tsc1* ^21^. Using this physiologically relevant system, we demonstrate that selective inactivation of SMG6 triggers robust innate and adaptive immune responses and potently suppresses liver tumorigenesis. Mechanistically, we uncover that SMG6 loss induces a type I interferon (IFN) response via activation of the cytosolic double-stranded RNA (dsRNA) sensor MDA5, revealing an unexpected physiological role for NMD in maintaining cytoplasmic dsRNA homeostasis and preventing aberrant activation of antiviral immunity. In support of the long-standing neoantigen hypothesis, we further find that the adaptive immune response is driven by the stabilisation and translation of NMD-targeted transcripts, which give rise to non-canonical peptides presented on MHC-I and elicit CD8^+^ T-cell infiltration. Together, our findings establish SMG6-dependent NMD as a central regulator of immune tolerance in the liver and highlight its selective inhibition as a promising strategy to promote tumour immunogenicity.

## Results

### SMG6 nuclease loss blocks tumour formation in an HCC model

We recently developed a conditional NMD loss-of-function mouse model that specifically targets the endonucleolytic activity of SMG6 (**Supplementary Fig. 1a)** ^20^. This is achieved through Cre/loxP-mediated conversion of a wild-type protein-expressing *Smg6^flox^* allele to *Smg6^mut^*, in which point mutations inactivate catalytic residues within the PIN nuclease domain. This approach overcomes the limitations of full NMD factor knockouts, which frequently impair cellular fitness due to both NMD-dependent and -independent functions of these proteins ^18, 19^. Liver-specific *Smg6^mut^* mice (i.e., *Smg6^flox/flox^*; *Alb^CreERT2^* treated with tamoxifen) exhibit only mild phenotypes ^20^, underscoring the suitability of this model for probing NMD function *in vivo*, including in disease-relevant settings.

To investigate the consequences of SMG6 inactivation on tumorigenesis, we combined this allele with a genetic HCC model (**Fig. 1a**). This model employs conditional knockout alleles of the tumour suppressors *Pten* and *Tsc1*, which encode negative regulators of the mTOR pathway. Liver-specific deletion of these genes (hereafter L-dKO) drives progression to non-alcoholic fatty liver disease (NAFLD), non-alcoholic steatohepatitis (NASH) and, ultimately, HCC ^21, 22^, closely recapitulating human disease evolution ^23^.

**Figure 1.**
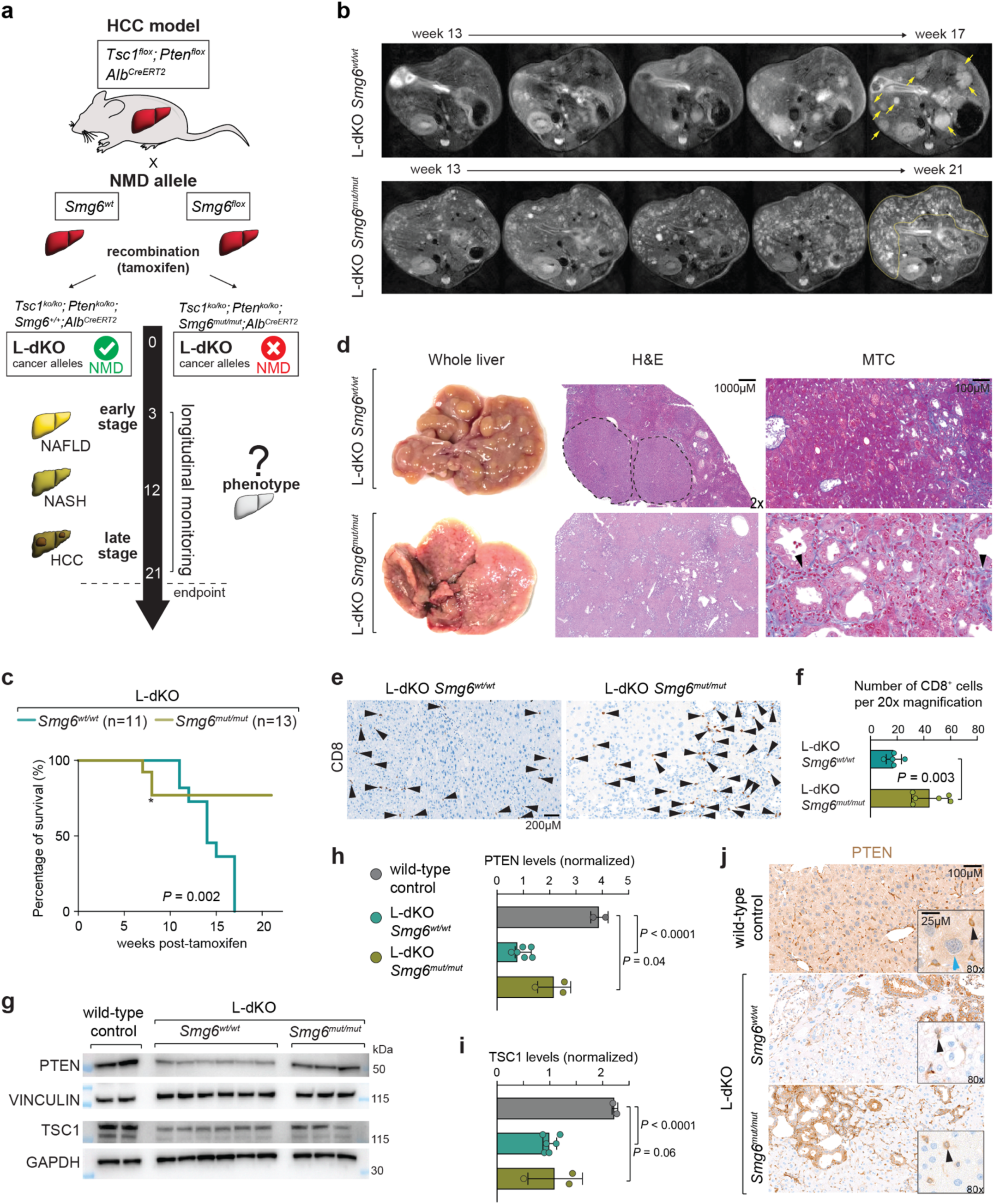
*Smg6* mutation suppresses cancer in genetic HCC model. (a) Schematic of the experimental design. Inactivation of tumour suppressor genes *Tsc1* and *Pten* via Cre-mediated recombination in liver (L-dKO) leads to tumours/HCC from week 12 ^21^. Cre-mediated recombination of the *Smg6^flox^* allele to *Smg6^mut^* introduces point mutations in catalytically important residues of the PIN domain ^20^. Both L-dKO *Smg6^wt/wt^* and *Smg6^flox/flox^*mice received 4 intraperitoneal tamoxifen injections to activate hepatocyte-specific CreERT2 (driven from *Albumin* promoter). Early-stage experiments were conducted 3 weeks post-tamoxifen. Late-stage animals were scanned via MRI from week 12 to 21 (endpoint). In L-dKO *Smg6^wt/wt^*, livers show NAFLD and NASH as indicated. (b) Representative longitudinal MRI scans for a tumour-bearing L-dKO *Smg6^wt^* mouse (top) and a tumour-free L-dKO *Smg6^mut^* animal. Tumours in L-dKO *Smg6^wt^*liver are indicated with yellow arrows (upper). Fibrotic plaques marked by yellow line in L-dKO *Smg6^mut^* liver (lower). (c) Kaplan-Meier plot, from two different cohorts, displays survival percentage of L-dKO *Smg6^wt^* (n = 11) and *Smg6^mut^* (n = 13) mice throughout the experiment. *P*-value by a log-rank (Mantel-Cox) test. Asterisk indicates 3 animals that deceased for other reasons than HCC in one of the cohorts (stress in animal handling). (d) Representative whole liver images, Masson’s Trichrome (MTC) staining for fibrosis (at 20x magnification) and H&E images (at 2x magnification) from late-stage L-dKO *Smg6^wt^* and *Smg6^mut^*mice. Dashed lines surround tumours in L-dKO *Smg6^wt^*. Arrowheads indicate fibrotic areas in L-dKO *Smg6^mut^*. (e) Representative immunohistochemistry (IHC) images of livers from late-stage mice, showing CD8^+^ cells, at 20x magnification. Arrowheads mark CD8^+^ T-cells (brown). (f) Quantification of CD8 IHC images. For each sample, the number of CD8^+^ T-cells was counted in 15 randomly selected images, at 20x magnification (L-dKO *Smg6^wt^*, n = 5 mice; L-dKO *Smg6^mut^*, n = 6 mice). The *P*-value was calculated with two-tailed unpaired *t*-test. (g) Western blot analysis of total liver proteins from late-stage mice for PTEN and TSC1. VINCULIN and GAPDH served as loading controls (Control, n = 2 mice; L-dKO *Smg6^wt^*, n = 6 mice; L-dKO *Smg6^mut^*, n = 3 mice). (h) Quantification of PTEN and VINCULIN western blots from (g) (i) Quantification of TSC1 and GAPDH western blots from (g). (j) Representative IHC images of livers from late-stage mice stained with anti-PTEN antibodies. Light brown staining shows PTEN expression in hepatocytes in wild-type control (blue arrowhead). Dark brown staining indicates PTEN expression in non-hepatocyte cells in both wild-type control and L-dKO (black arrowhead). Images were captured at 20x (large images) and 80x (small images) magnifications. For (f), (h), and (i) data are plotted as means, and the error bars indicate SEM. The *P*-values were calculated with two-tailed unpaired *t*-test.

We first assessed the impact of *Smg6^mut^* on HCC formation at late stages (starting week 12 post-tamoxifen). Weekly magnetic resonance imaging (MRI) revealed rapidly progressing liver tumours in L-dKO *Smg6^wt^* mice (**Fig. 1b**), necessitating euthanasia by week 17 (**Fig. 1c**). In contrast, L-dKO *Smg6^mut^* mice developed no tumours, and none required sacrifice, indicating complete suppression of HCC.

Macroscopic and histological analyses confirmed HCC in L-dKO *Smg6^wt^* (**Fig. 1d**, top). L-dKO *Smg6^mut^*mice, which persisted until the week 21 endpoint, displayed prominent duct proliferation and moderate perisinusoidal and periportal fibrosis (**Fig. 1d**, bottom), leading to a nodular parenchymal architecture but no histological evidence of HCC. Notably, L-dKO *Smg6^mut^* livers showed elevated CD8^+^ T-cell infiltration relative to L-dKO *Smg6^wt^*(**Fig. 1e, f**), consistent with sustained adaptive immune activation. Serum liver-injury markers (ALT, AST, LDH; **Supplementary Fig. 1b-d**) and hepatomegaly (**Supplementary Fig. 1e**) were similarly elevated in both groups. As essential controls, L-dKO *Smg6^mut^* livers retained expression of recombined *Smg6^mut^* mRNA (**Supplementary Fig. 1f**) and lacked PTEN and TSC1 specifically in hepatocytes (**Figures 1g-i**), with residual protein restricted to non-parenchymal cells such as Kupffer cells (**Figure 1j**). Additional histological analysis revealed that the more pronounced duct proliferation in L-dKO *Smg6^mut^* livers was largely composed of biliary epithelial cells (i.e., non-hepatocyte), indicating that higher remaining PTEN signal in the L-dKO mutant livers reflects altered tissue composition rather than escape from recombination (**Supplementary Fig. 2a, b**). Together, these findings rule out trivial explanations – such as synthetic lethality of the *Smg6^mut^*; *Pten^ko^*; *Tsc1^ko^* genotype or compensatory outgrowth of non-mutant cells – and demonstrate that SMG6 nuclease inactivation robustly prevents tumour development in this genetic model of HCC.

### SMG6 inactivation induces early immune infiltration

Given the striking phenotypic divergence at late stages – tumours in *Smg6* wild-type mice but none in *Smg6* mutants – we next investigated earlier time-points in our model, focusing on 3 weeks post-tamoxifen (**Fig. 1a**), to understand the molecular and cellular changes occurring in NMD-deficient livers prior to tumour development. As expected, western blot analysis at this stage confirmed strong depletion of TSC1 and PTEN proteins in L-dKO livers regardless of *Smg6* genotype (**Fig. 2a, b; Supplementary Fig. 1j-l**).

**Figure 2.**
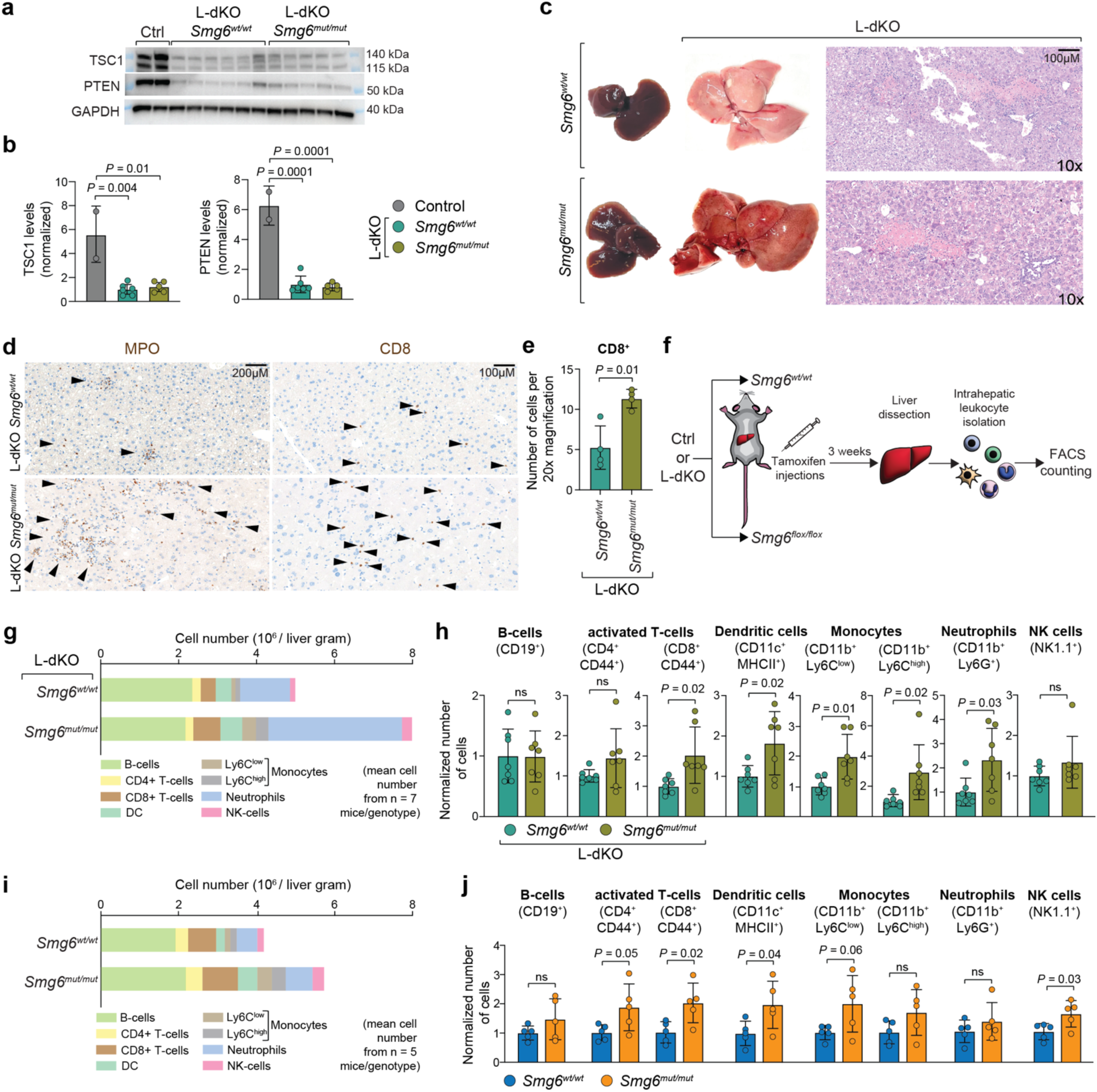
SMG6-dependent early inflammation. (a) Western blot analysis on total liver proteins, for PTEN and TSC1, at early-stage post-tamoxifen. GAPDH serves as loading control (Control, n = 2 mice; L-dKO *Smg6^wt^*, n = 6 mice; L-dKO *Smg6^mut^*, n = 5 mice). (b) Quantification of western blots from (a). (c) Representative images of whole-livers from early-stage *Smg6^wt^* and *Smg6^mut^*mice, without and with L-dKO background, as well as Hematoxylin and Eosin (H&E)-stained sections (at 10x magnification) from early-stage L-dKO *Smg6^wt^* and *Smg6^mut^*mice. (d) Representative IHC images of livers from early-stage L-dKO *Smg6^wt^* and *Smg6^mut^* mice, using neutrophil marker Myeloperoxidase (MPO) and T-cell marker CD8, at 20x magnification. Arrowheads mark MPO-positive or CD8^+^ cells. (e) Quantification of CD8-stained IHC images. For each sample, the number of CD8^+^ T-cells was counted in 15 randomly selected images, at 20x magnification (mean values from 4 mice per group). (f) Schematic of the experimental design for intrahepatic leukocyte isolation and flow cytometry-based counting. *Smg6^wt/wt^*and *Smg6^flox/flox^* mice (with or without L-dKO) received 5 intraperitoneal tamoxifen injections. Livers were dissected 3 weeks later, and leukocytes were isolated for flow cytometric quantification. FACS, fluorescence-associated cell sorting. (g) Overall leukocyte numbers per gram liver tissue (mean values from 7 mice per group), as assessed by flow cytometry. Cells are identified according to indicated color code. DC, dendritic cells; NK-cells, natural killer cells. (h) Relative leukocyte counts for individual cell types as assessed by flow cytometry. Graphs show normalized numbers (set to 1 in L-dKO *Smg6^wt/wt^*) of indicated leukocyte type (n = 7 mice for each group). (i) Overall leukocyte numbers per gram liver tissue (mean values from 5 mice per group), as assessed by flow cytometry. Cells are identified according to indicated color code. (j) Relative leukocyte counts for individual cell types as assessed by flow cytometry. Graphs show normalized numbers (set to 1 in *Smg6^wt/wt^*) of indicated leukocyte type (n = 5 mice for each group. Data are plotted as means, and the error bars indicate SEM. For (b), (e), (h), and (j) data are plotted as means, and the error bars indicate SEM. The *P*-values were calculated with two-tailed unpaired *t*-test.

Macroscopic and histological analyses revealed comparable NASH-like features in L-dKO *Smg6^wt^* and *Smg6^mut^* livers, including focal microsteatosis, hepatocyte ballooning, inflammatory infiltration, and necrosis (**Fig. 2c; Supplementary Fig. 3a**). However, inflammation was already markedly more pronounced in L-dKO *Smg6^mut^* livers, as shown by increased staining for the neutrophil/monocyte marker myeloperoxidase (MPO; **Fig. 2d**, left panels) and the cytotoxic T-cell marker CD8 (**Fig. 2d**, right panels; **Fig. 2e**). Flow cytometry further confirmed elevated hepatic leukocyte counts in L-dKO *Smg6^mut^*livers (**Fig. 2f-h; Supplementary Fig. 3b, c**). Consistent with this heightened inflammatory state, serum liver-injury markers peaked specifically in L-dKO *Smg6^mut^*animals at this early time point (**Supplementary Fig. 1b-d**).

Interestingly, increased immune cell infiltration was also observed in *Smg6^mut^* livers lacking the L-dKO alleles, indicating that NMD inhibition alone promotes a pro-inflammatory hepatic environment (**Fig. 2i, j**). Notably, when *Smg6* mutation was combined with L-dKO alleles, infiltration was further exacerbated – particularly among innate immune populations such as neutrophils and Ly6C^high/low^ monocytes (**Fig. 2g-j**). In addition, L-dKO *Smg6^mut^*livers showed elevated apoptotic markers (**Supplementary Fig. 3d-h**) and increased expression of pro-fibrotic genes (**Supplementary Fig. 3i**), consistent with enhanced cell death and liver injury. Together, these findings demonstrate that SMG6 inactivation potentiates early liver inflammation, immune cell infiltration, and apoptosis in a tumour-prone background – well before overt HCC formation in this model.

### SMG6 inactivation induces CD8⁺ T-cell responses through presentation of non-canonical immunopeptides

To investigate the mechanism underlying the immune-altered state in *Smg6^mut^* livers, we first focused on adaptive immune activation – particularly the strong infiltration of CD8^+^ T-cells (**Fig. 2j**), key effectors of anti-tumour immunity. We hypothesised that NMD inhibition enables hepatocytes to present novel MHC-I-bound immunopeptides derived from stabilised NMD-target transcripts.

To test this, we employed a multi-omics strategy (**Fig. 3a**) combining RNA-seq (to identify NMD-regulated transcripts), ribosome profiling (Ribo-seq; to detect translated open reading frames), and immunopeptidomics (mass spectroscopy-based identification of MHC-I-bound peptides). This integrative approach allowed us to identify peptides uniquely presented in *Smg6^mut^*livers and trace them back to their transcriptomic and translational origins.

**Figure 3.**
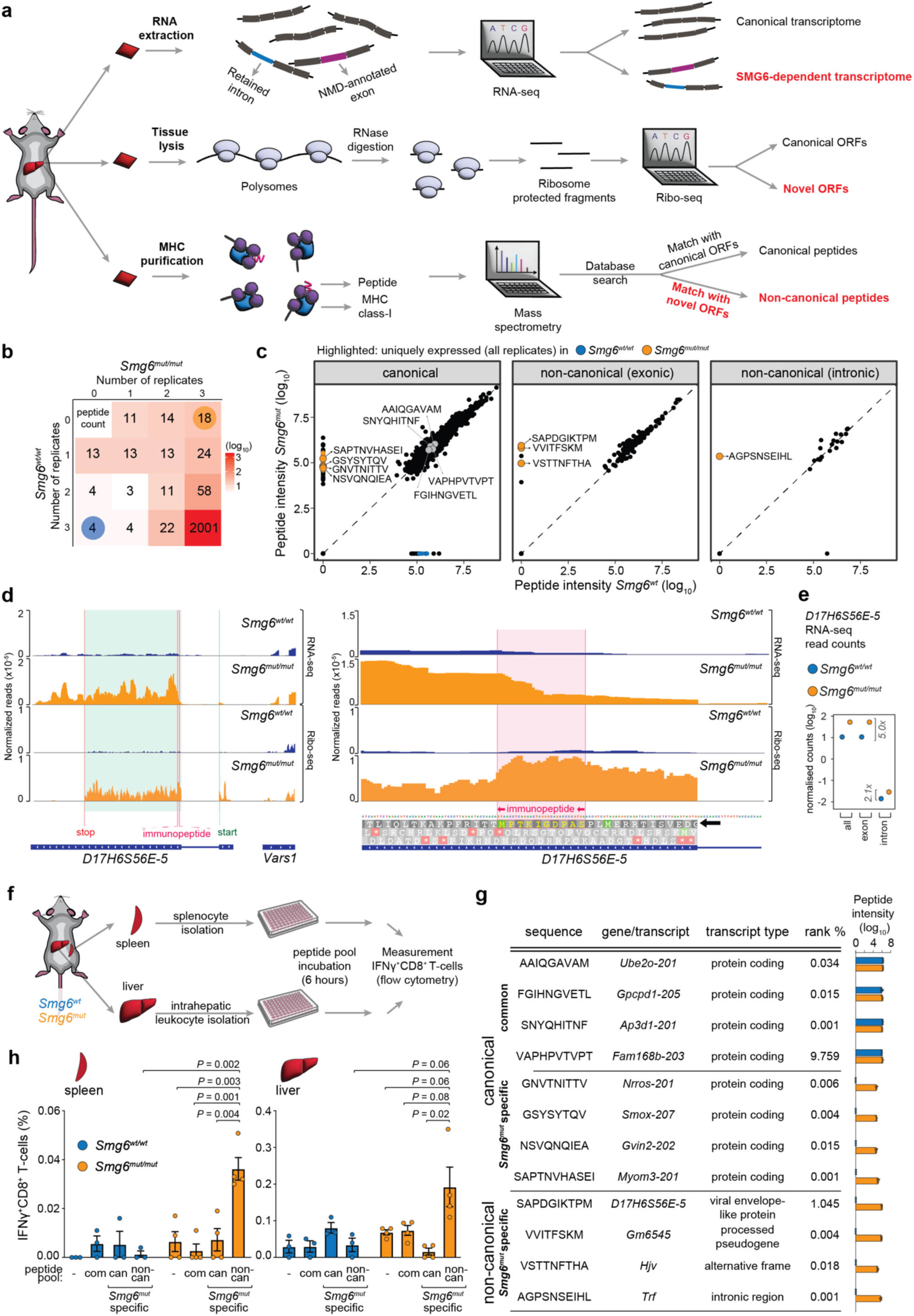
Multiomics-based identifcation and validation of SMG6-dependent immunopeptides. (a) Schematic of the multiomics approach used to identify SMG6-dependent immunopeptide candidates by combining RNA-seq, Ribosome profiling (Ribo-seq) and immunopeptidomics data. (b) MHC-I binder peptide count and distribution across *Smg6^wt/wt^* and *Smg6^mut/mut^*liver replicates (n = 3 mice per genotype). Enrichment of unique peptides in *Smg6^mut/mut^* samples (18 in all replicates, orange circle) as compared to *Smg6^wt/wt^* (4 in all replicates, blue circle) is highly significant (*P* = 0.000053; Fisher’s exact test). The heatmap scale is colored according to peptide count, for better visual representation. (c) Scatter plots of peptide intensity values, showing peptide categories canonical (protein-coding), non-canonical (exonic) and non-canonical (intronic), in *Smg6^wt/wt^*(x-axis) vs, *Smg6^mut/mut^* (y-axis). Data are plotted as medians from the three replicates. Unique peptides for one of the genotypes that are found in all three replicates are colored orange (*Smg6^mut/mut^* -specific peptides) and blue (*Smg6^wt/wt^*-specific peptides), with specific peptides that are used in the downstream pools highlighted and annotated with their amino acid sequence. (d) RNA-seq (top panels) and Ribo-seq (bottom panels) read pileup plots show the strongly increased transcript and footprint abundance of *D17H6S56E-5* in *Smg6^mut/mut^*. Left panel shows the larger genomic region, including the CDS (highlighted in turquoise) from which the non-canonical polypeptide is translated, while the right panel zooms into the specific region where the immunopeptide is translated (highlighted in light magenta). Plots were generated using merged sequencing data from the three biological replicates (individual livers) per group. Please note gene orientation from right to left. (e) Normalized exonic and intronic RNA-seq read counts of *D17H6S56E-5*. Each dot represents the merged sequencing data from three biological replicates/livers. (f) Schematic of the immunogenicity quantification experiment. *Smg6^wt/wt^* and *Smg6^mut/mut^* mice were sacrificed 3 weeks post-tamoxifen. Spleens and livers were extracted from the animals and leukocytes were isolated. The cells were then incubated with peptide pools indicated in (g), for 6 hours. The percentages of IFNγ^+^CD8^+^ T-cells were used as a readout for the immunogenicity of each peptide pool. Common peptides were used as negative control (*Smg6^wt/wt^*, n = 3 mice, *Smg6^mut/mut^*, n = 4 mice). (g) Table showing peptide sequences derived from the indicated genes, transcript types, rank percentages from NetMHCpan prediction and their intensity in each of three replicates of *Smg6^wt/wt^*and *Smg6^mut/mut^* livers. Each group of four peptides was used as a pool in the experiment in (h). (h) Overall IFNγ^+^CD8^+^ T-cell percentage per tissue (left panel, spleen; right panel, liver), as assessed by flow cytometry, upon treatment with no peptide pool (-), canonical common peptide pool (com), canonical *Smg6^mut^*-specific peptide pool (can), and non-canonical *Smg6^mut^*-specific peptide pool (non-can). Bars indicate means, and the error bars SEM; individual datapoints are measurements from independent animals (*Smg6^wt/wt^*, n = 3 mice, *Smg6^mut/mut^*, n = 4 mice). The *P*-values were calculated with two-tailed unpaired *t*-test.

Immunopeptidomics analyses identified over 2000 MHC-I-bound peptides (**Fig. 3b**), the majority shared between *Smg6^mut^* and *Smg6^wt^* livers and mapping to canonical coding regions (**Fig. 3c; Supplementary Table 1**). However, 18 peptides were consistently detected exclusively in *Smg6^mut^* livers across all replicates, compared to only four *Smg6^wt^*-specific peptides – representing a highly significant enrichment of *Smg6^mut^*-specific antigens (*P* = 5.4e-05; Fisher’s exact test). Four of these peptides originated from non-canonical coding regions: three from exonic sequences and one from an intronic region (**Fig. 3c**).

For example, the peptide SAPDGIKTPM mapped to the *D17H6S56E-5* gene, previously annotated as a long non-coding RNA with homology to a mouse leukaemia virus envelope gene ^24^. RNA-seq analysis revealed markedly increased exon-derived reads in *Smg6^mut^* livers (**Fig. 3d**), whereas intronic reads showed only modest elevation (**Fig. 3e**), suggesting post-transcriptional stabilisation consistent with an NMD substrate. Ribo-seq confirmed translation of a 596-aa ORF containing the SAPDGIKTPM epitope near its N-terminus (**Fig. 3d**, right). Similarly, the VVITFSKM peptide originated from the *Gm6545*/*A330040F15Rik* pseudogene locus (a paralogue of *Fam111a*), which showed increased RNA expression and ribosome occupancy of a 554-aa ORF (**Supplementary Fig. 4a**). Two additional peptides – VSTTNFTHA and AGPSNSEIHL – mapped to an out-of-frame transcript from the *Hjv* locus and to an intronic region of *Trf*, respectively (**Supplementary Table 1**). Together, these findings indicate that NMD inhibition enables the expression and translation of non-canonical transcripts, which give rise to novel MHC-I-presented peptides, potentially explaining the observed CD8^+^ T-cell activation.

We next assessed whether these peptides were immunogenic *in vivo*. All four candidates exhibited strong predicted MHC-I binding affinity (NetMHCpan rank < 2%; **Fig. 3g**). We chemically synthesised the four non-canonical peptides and tested them as a pool on splenic and hepatic leukocytes isolated three weeks post-tamoxifen to determine immunogenic potential (**Fig. 3f**). To assess specificity, we compared this pool with two control peptide pools: one composed of commonly detected peptides and one consisting of *Smg6^mut^*-specific canonical peptides (**Fig. 3c, g**). Upon stimulation with the *Smg6^mut^*-specific non-canonical peptide pool, CD8^+^ T-cells from both spleen and liver of *Smg6^mut^* mice showed a robust increase in IFN-γ expression (**Fig. 3h; Supplementary Fig. 4b, c**). In contrast, leukocytes from *Smg6^wt^*mice showed no such response, and neither control pool elicited IFNγ^+^CD8^+^ T-cell activation in either genotype (**Fig. 3h**). All non-canonical peptides individually also induced IFN-γ expression in splenic and hepatic CD8^+^ T-cells of *Smg6^mut^* mice (**Supplementary Fig. 5a-d**). Together, these results demonstrate that *Smg6^mut^*-specific non-canonical peptides are processed and presented via MHC-I and are sufficient to elicit robust CD8^+^ T-cell activation *in vivo*.

### SMG6 inactivation triggers a cell-autonomous type I IFN response

We next investigated whether the observed innate immune infiltration observed in *Smg6^mut^* livers might arise from a cell-intrinsic, SMG6-dependent mechanism. Supporting this possibility, RNA-seq analysis revealed broad upregulation of transcripts indicative of an activated type I IFN response (**Fig. 4a**) ^25^. Quantitative real-time PCR (qPCR) confirmed strong *Smg6^mut^*-dependent induction of canonical IFN-stimulated genes (ISGs), including *Ifih1*, *Oas1a*, *Ifi44*, *Zbp1*, and *Ifit1* (**Fig. 4b**), consistent with robust activation of the type I IFN pathway.

**Figure 4.**
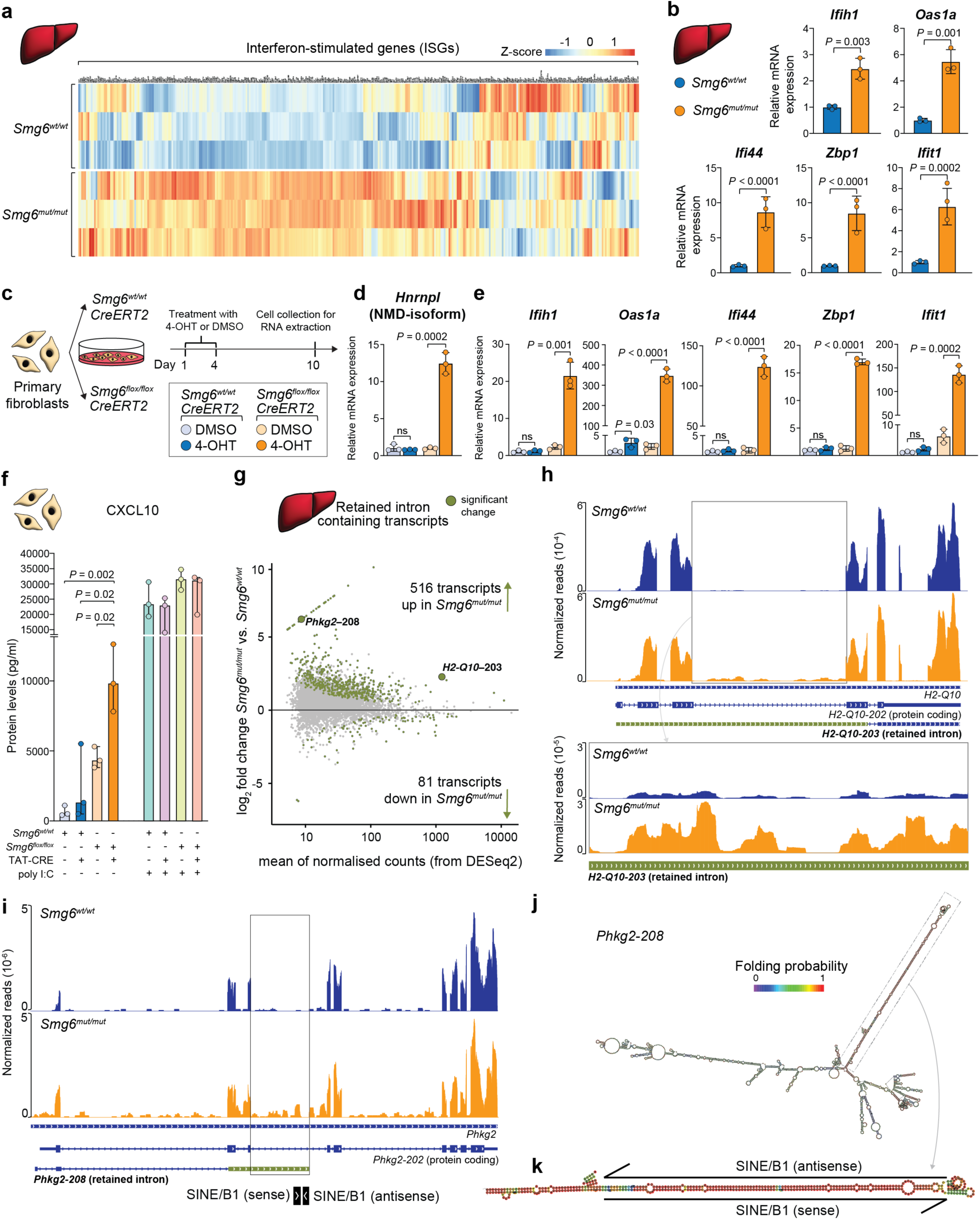
*Smg6* mutation elicits type I IFN response and increased intron abundance. (a) Heatmap showing ISG expression in control and mutant livers (n = 3 livers per group). RNA-seq data was plotted after Z-score transformation of normalized expression levels across the 6 samples. (b) Relative hepatic mRNA levels of indicated ISGs (n = 3 mice for each group), detected via qPCR. *Smg6^wt/wt^*(control) samples were set to 1. (c) Schematic of the experiment to analyze SMG6-dependent ISG expression. To induce *Smg6* mutation in *Smg6^flox/flox^*; *CreERT2* fibroblasts, cells were treated with 2 µM 4-hydroxytamoxifen (4-OHT) during days 1 to 4, and cells were collected 6 days later (day 10) for RNA isolation and qPCR. Control treatment was DMSO. (d) Relative mRNA expression of the endogenous, diagnostic NMD-target, an *Hnrnpl* splice form, in samples collected as described in (c). Three biological replicates were used per group. (e) Relative mRNA expression of indicated ISGs in same samples as in (d). Three biological replicates were used per group. (f) ELISA Analysis of CXCL10. At day 1, *Smg6^wt/wt^* and *Smg6^flox/flox^* cells were treated with TAT-CRE (2.5 µM, 8 hours) or vehicle. At day 7, indicated groups were treated with poly I:C to serve as positive control (1 µM, 8 hours). At day 10, supernatants were collected, and CXCL10 levels were detected via ELISA. Data are plotted as medians (n = 3 biological replicates for each group). The *P*-values were calculated with two-tailed unpaired *t*-test. (g) Differential expression analysis at the transcript level, specifically on retained intron containing mRNAs, comparing *Smg6^wt/wt^* and *Smg6^mut/mut^* livers (n = 3 mice per genotype). Significantly more or less abundant transcripts are indicated in olive, showing strong skew to upregulation. Two transcripts are individually labeled, as they are further shown in panels (h) and (i) (h) RNA-seq pileup plots show the upregulated transcript isoform (containing intron) for *H2-Q10*. Top panels show the larger genomic region, including the retained intron, while bottom panels zoom in the retained intron area. Plots were generated using merged sequencing data from three biological replicates per group. Please note gene orientation from left to right. (i) RNA-seq pileup plots as in (h) show the increased abundance of the intron-containing transcript isoform for *Phkg2.* Genomic area where SINE/B1 elements are located in intronic region is highlighted. Plots were generated using merged sequencing data from three biological replicates per group. Please note gene orientation from left to right. (j) RNA structure prediction of *Phkg2-208* by RNAfold web server, showing extensive predicted dsRNA region. (k) Predicted double-stranded structure from SINE/B1 elements located in the intronic region of *Phkg2-208* transcript. For (b), (d) and (e) data are plotted as means, and the error bars indicate SEM. The *P*-values were calculated with two-tailed unpaired *t*-test.

Because whole-liver tissue contains a mixture of *Smg6^mut^*hepatocytes and *Smg6^wt^* immune cells, ISG induction could reflect either hepatocyte-leukocyte interactions or a cell-autonomous consequence of SMG6 loss. To distinguish between these possibilities, we used *Smg6^flox/flox^*dermal tail fibroblasts transduced with tamoxifen-activatable *CreERT2* (**Fig. 4c**) ^20^. Tamoxifen treatment efficiently abrogated NMD, as evidenced by a >10-fold increase in an NMD-sensitive *Hnrnpl* splice isoform (**Fig. 4d**). Importantly, NMD inhibition triggered massive ISG upregulation, with *Ifih1*, *Oas1a*, *Ifi44*, *Zbp1*, and *Ifit1* increasing by up to several hundred-fold (**Fig. 4e**), establishing that loss of SMG6 activity is sufficient to induce a type I IFN response in a cell-autonomous manner.

To determine whether this type I IFN response could be recapitulated by other approaches that target NMD, we inhibited the UPF1-specific kinase SMG1 using a small-molecule inhibitor (SMG1i) ^26^ in *Smg6^flox^* fibroblasts and compared ISG induction with that in *Smg6^mut^* cells (**Supplementary Fig. 6a**). 24-hour SMG1i treatment robustly increased the NMD-sensitive *Hnrnpl* isoform (**Supplementary Fig. 6b**), confirming effective NMD impairment. ISG induction was clearly detectable, though quantitatively weaker than in *Smg6^mut^* cells (**Supplementary Fig. 6c**). However, because SMG1i rapidly causes toxicity and is known to inhibit additional PI3K-like kinases, prolonged inhibition as in the *Smg6^mut^* model – likely required to establish a fully developed type I IFN response – was not feasible. Thus, these data are consistent with ISG activation being a general consequence of NMD inhibition, yet the magnitude of the response cannot be directly compared between SMG1i-treated and *Smg6^mut^* cells.

To assess IFN pathway activation at the protein level, we measured CXCL10, an IFN-inducible chemokine and well-established surrogate for type I IFN signalling ^27^. CRE-treated *Smg6^flox^* fibroblasts displayed markedly elevated CXCL10 levels relative to all relevant controls (**Fig. 4f**), further supporting autonomous type I IFN pathway activation under *Smg6^mut^* conditions. Importantly, all genotypes responded comparably to poly(I:C) stimulation, reaching similarly high CXCL10 levels (**Fig. 4f**), indicating that the downstream IFN signalling machinery remains intact and fully inducible. Together, these findings demonstrate that SMG6 endonuclease inactivation elicits a cell-intrinsic type I IFN response in both liver tissue and fibroblasts. This response can occur independently of immune cell interactions,.

### SMG6 inactivation increases the abundance of intron-retaining transcripts in liver

The type I IFN response is a key antiviral pathway triggered when cytoplasmic sensors detect atypical nucleic acids, such as dsRNA. Although typically associated with viral infection, endogenous dsRNA – arising from dysregulated RNA processing – can similarly enhance anti-tumour immunity or induce apoptosis. One established source of endogenous dsRNA is splicing inhibition, which promotes cytoplasmic leakage of intron-containing mRNAs. These transcripts can form dsRNA structures, often due to repetitive element sequences embedded within introns ^28–30^.

Transcripts with retained introns are well-established NMD substrates ^31, 32^, as also shown in our previous work using *Smg6^mut^* fibroblasts ^20^. We therefore asked whether a similar accumulation occurs in *Smg6^mut^*livers. Differential expression analysis of hepatic RNA-seq data focusing on annotated intron-retaining isoforms revealed a marked increase in over 500 such isoforms in *Smg6^mut^* livers, compared to only 81 showing significant downregulation (**Fig. 4g**). Inspection of representative loci confirmed that exonic read coverage – shared by canonical and intron-retaining isoforms – remained largely unchanged, while intronic read coverage was markedly elevated (**Fig. 4h, i**), supporting specific enrichment of intron-containing transcripts. Notably, several retained introns were predicted to form dsRNA structures due to oppositely oriented repetitive elements. For example, the retained intron in *Phkg2-208* (**Fig. 4i**) contains two SINE/B1 elements in sense and antisense orientation, generating extensive complementarity predicted to support intramolecular dsRNA formation (**Fig. 4j, k**). Additional transcript category analyses revealed a global increase in lncRNAs and annotated NMD targets, alongside a reduction in protein-coding isoforms (**Supplementary Fig. 6d-f**).

Together, these findings demonstrate that SMG6 inactivation leads to widespread accumulation of intron-retaining transcripts in the liver, providing plausible endogenous sources of immunostimulatory dsRNA that may contribute to activation of the type I IFN pathway.

### MDA5 is required for the SMG6-dependent type I IFN response

The widespread accumulation of intron-retaining transcripts in *Smg6^mut^*livers and fibroblasts suggests a source of endogenous dsRNA capable of triggering innate immune signalling. To identify how this immunogenic RNA is sensed, we considered the cytoplasmic dsRNA receptors MDA5 and RIG-I, both of which activate type I IFN responses through MAVS ^33^. Since RIG-I preferentially recognises short dsRNAs with 5′-triphosphate ends – a hallmark of viral RNAs ^33^ – we hypothesised that MDA5, which detects longer, structured dsRNA species ^34^, mediates the response to *Smg6* inactivation.

To test this, we generated *Smg6^flox/flox^* fibroblast lines lacking *Ifih1*, the gene encoding MDA5 (**Fig. 5a**, **Supplementary Fig. 7a, b**), and induced *Smg6* mutation using recombinant, cell-permeable TAT-Cre protein (**Fig. 5b**). NMD inhibition remained intact in MDA5-deficient cells, as evidenced by increased accumulation of the NMD-sensitive *Hnrnpl* isoform (**Fig. 5c**). Strikingly, however, ISG induction – robust in MDA5-proficient *Smg6^mut^* cells – was largely abolished in the absence of MDA5 (**Fig. 5d**), demonstrating that this sensor is essential for the SMG6-dependent IFN response. Consistent with the accumulation of immunogenic RNA, cytoplasmic dsRNA levels were significantly elevated in *Smg6^mut^*fibroblasts, as detected by increased J2 antibody staining (**Fig. 5e, f**) ^35^.

**Figure 5.**
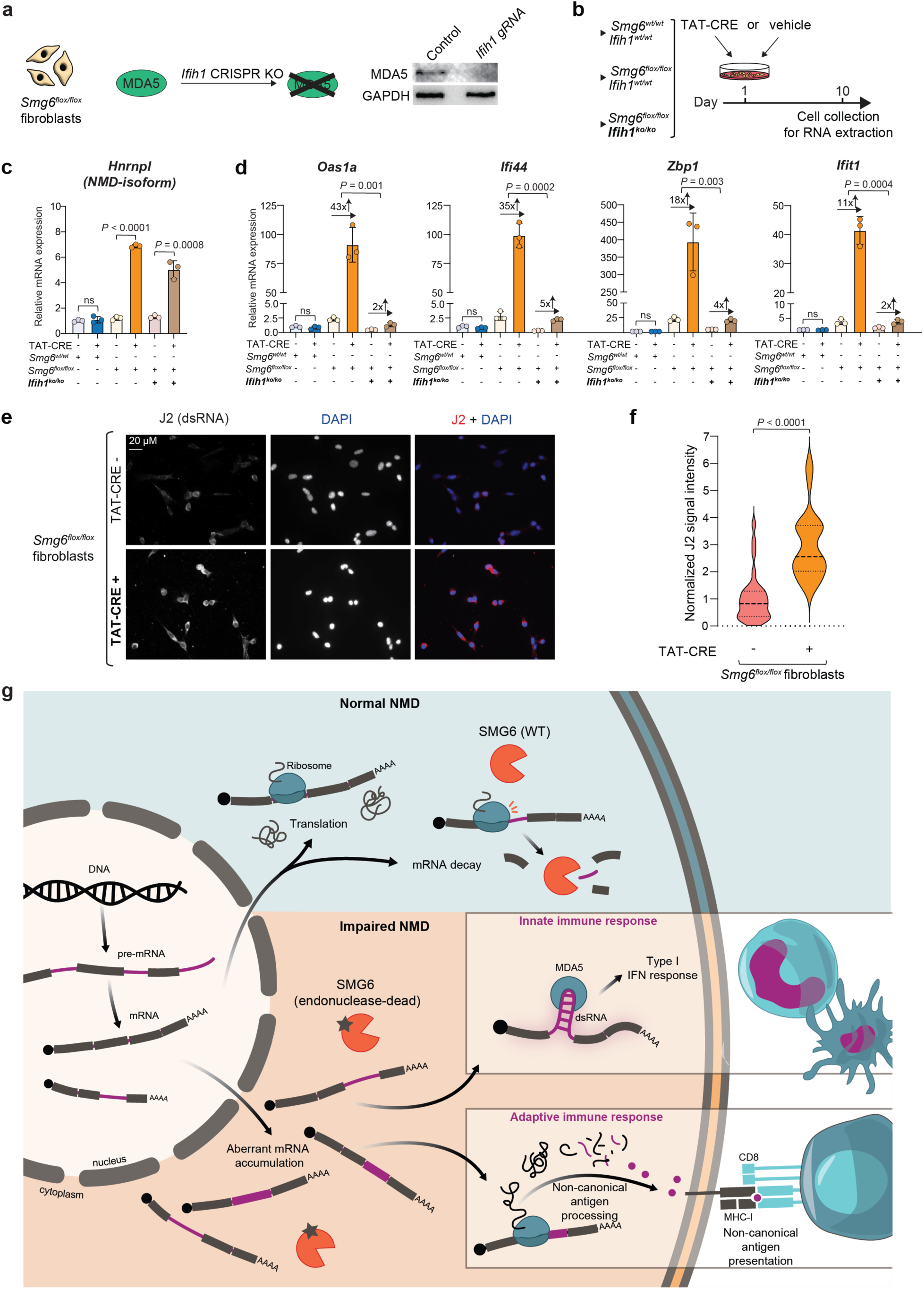
*Smg6* mutation drives type I IFN response through dsRNA sensor MDA5. (a) Western blot validation of the loss of MDA5 expression in *Ifih1^ko/ko^* fibroblasts (in background of *Smg6^flox/flox^* fibroblasts). GAPDH serves as loading control. (b) Schematic of the experiment. Cells with indicated genotypes were treated with either TAT-CRE (2.5 µM) or vehicle for 8 hours and then harvested at day 10 for RNA extraction and qPCR analyses. (c) Relative mRNA expression of the endogenous, diagnostic NMD-target, an *Hnrnpl* splice form, in samples collected as described in (b). Three biological replicates were used per group. (d) Relative mRNA expression of indicated ISGs. Arrows indicate the fold-change in the expression of indicated genes between cells nontreated vs. treated with TAT-CRE. Three biological replicates were used per group. (e) Representative immunofluorescence images using dsRNA-specific antibody J2 (red) on *Smg6^flox/flox^*fibroblasts that were not treated (top) or treated (bottom) with cell-permeable TAT-CRE to induce *Smg6* mutation. DAPI (blue) serves to visualize nuclei. (f) Quantification of J2 signal intensity (n = 238 cells for TAT-CRE-negative, n = 185 cells for TAT-CRE-positive). Median and third quartiles of each group are marked by dotted lines. The *P*-value was calculated with two-tailed unpaired *t*-test. (g) Model for how the suppression of NMD activity via endonuclease-inactive SMG6 activates both innate and adaptive branches of the immune system. For innate responses, aberrant mRNAs (shown here: intron-retained transcripts known to contain repetitive elements) can form double-stranded configurations, causing activation of cytoplasmic dsRNA sensor MDA5, which upregulates a type I IFN response, resulting in innate immune cell infiltration. With the accumulation of aberrant or otherwise NMD-suppressed mRNAs in the cytoplasm, their translation leads to the production of unusual/non-canonical peptides which are then subsequently processed for MHC-I presentation on the cell surface, which attracts CD8^+^ T-cells to infiltrate the tissue, leading to adaptive immune cell recognition. Please note that for simplicity, prior steps required for CD8^+^ T-cell activation (i.e., antigen presentation by dendritic cells) have been omitted from the schematic.

These results identify MDA5 as the critical sensor linking SMG6 inactivation to type I IFN activation through detection of endogenous dsRNA. Moreover, the specificity of this response – and its absence in Cre-treated MDA5-deficient cells – rules out confounding activation of alternative pathways such as cGAS-STING, which has been associated with Cre-induced DNA damage ^36^. Together, our findings establish SMG6-dependent NMD as a key RNA quality control mechanism that prevents the accumulation of endogenous dsRNA and inappropriate activation of MDA5 and type I IFN signalling, thereby preserving innate immune quiescence in non-infected cells.

### SMG6 deficiency slows tumour progression in an orthotopic HCC model

Because *Smg6* mutation completely prevented tumour development in our genetic L-dKO HCC model (**Fig. 1b-d**), this system did not allow us to assess how loss of SMG6 activity affects the growth of established tumours. To determine whether the tumour-suppressive effects of SMG6 inactivation extended to an independent setting, we turned to an orthotopic transplantation model. We generated *Smg6^ko/ko^* Hepa1-6 tumour cells (**Supplementary Fig. 8a, b**), a cell line originally derived from a C57BL/6 hepatoma ^37^. *Smg6^ko^* cells displayed similar cell cycle phase distribution compared to the *Smg6^wt^* parent cells (**Supplementary Fig. 8c**), in line with indistinguishable growth behaviour in cell culture. As expected, NMD inhibition and ISG upregulation were apparent in the *Smg6^ko^* line (**Supplementary Fig. 8d**), along with markedly elevated IFN-α/β levels (**Supplementary Fig. 8e**). To assess tumour progression *in vivo*, we orthotopically injected *Smg6^wt^* or *Smg6^ko^* Hepa1-6 cells into immunocompetent C57BL/6 mice and monitored tumour growth longitudinally by MRI (**Supplementary Fig. 9a**). During the first week after implantation, *Smg6^wt^* and *Smg6^ko^* liver tumours expanded at similar rates, indicating that *Smg6*-deficient cells were not rejected by the host immune system. Strikingly, during weeks 2 and 3, *Smg6^ko^* tumours exhibited significantly slower progression than their wild-type counterparts (**Supplementary Fig. 9b-e**). Histopathological analyses of these tissues showed increased CD4^+^ and CD8^+^ T-cell infiltration in *Smg6^ko/ko^* tumours compared to the wild-type tumours (**Supplementary Fig. 9f-i**). Together, these findings demonstrate that *Smg6*-deficient tumour cells give rise to tumours with impaired outgrowth in an orthotopic HCC model. The reduced progression of *Smg6*-deficient tumours cannot be attributed to intrinsic proliferation defects of the tumour cells and therefore supports a role for SMG6 in constraining anti-tumour immunity across distinct experimental systems.

## Discussion

Our study reveals that SMG6-dependent NMD plays a dual and previously underappreciated role in shaping immune homeostasis and tumour development in the liver. Using a hepatocyte-specific, immunocompetent genetic HCC model, we show that selective inactivation of the SMG6 endonuclease elicits both innate and adaptive immune responses and completely suppresses tumour formation (**Fig. 5g**).

At the innate immune level, we uncover an unexpected innate mechanism: SMG6 prevents the cytoplasmic accumulation of endogenous dsRNAs for which intron-retaining transcripts represent a likely source. Upon SMG6 inactivation, these RNAs accumulate, engage the cytosolic sensor MDA5, and trigger a potent type I IFN response. MDA5 deletion abolishes ISG induction while preserving NMD inhibition, establishing endogenous dsRNA sensing as the mechanistic axis connecting NMD disruption to innate immune activation. Thus, SMG6 acts as a molecular gatekeeper that maintains immune quiescence by preventing aberrant dsRNA accumulation.

In parallel, at the adaptive immune level, our multi-omics analyses demonstrate that transcripts normally eliminated by NMD give rise to non-canonical MHC-I-presented peptides capable of activating CD8^+^ T cells. These data provide *in vivo* validation of the long-standing “neoantigen hypothesis”, in which NMD inhibition exposes immune-relevant epitopes. Our immunopeptidomics analyses identified four high-confidence non-canonical immunopeptides uniquely presented in *Smg6^mut^*livers. While limited in number, these peptides were identified under stringent criteria requiring reproducible detection in all replicates, strong predicted MHC-I binding, and direct Ribo-seq support. Each peptide independently activated CD8^+^ T cells, and their appearance as early as three weeks after induction aligns with the timeframe when immune surveillance is effective. Future *in vivo* functional intervention studies, including CD8^+^ T-cell depletion ^38^ or IFNAR blockade ^39^, will be important to establish the relative contributions of adaptive and innate responses to tumour suppression.

Together, these findings establish a coordinated model in which SMG6 simultaneously restricts endogenous dsRNA-mediated innate immune activation and limits the generation of non-canonical immunopeptides, thereby maintaining immune tolerance under physiological conditions.

This dual function reframes the conceptual understanding of NMD. Historically viewed as a system preventing translation of truncated proteins, NMD has been interpreted mainly through a protein-centric lens. Our findings add an equally important RNA-centred dimension: NMD safeguards the cytoplasm from endogenous dsRNAs that would otherwise activate antiviral signalling. In this respect, SMG6 parallels other dsRNA-modifying or -processing factors such as ADAR1, which prevent inappropriate activation of innate immune sensors and maintain self-tolerance ^40^. Viewed from an evolutionary perspective, this RNA-centric function may represent a key selective pressure driving the conservation of the pathway. While suppression of toxic proteins remains a valid functional rationale, the more immediate threat posed by the activation of innate immune sensors – such as MDA5 and PKR – by endogenous RNAs may have exerted particularly strong evolutionary pressure. These sensors are exquisitely tuned to recognise features common to viral nucleic acids but can be inadvertently activated by endogenous transcripts when RNA processing or degradation fails. In this context, SMG6-mediated degradation acts as a buffer, maintaining the cytoplasmic RNA environment below an immunostimulatory threshold. The parallels to mechanisms implicated in autoimmune and inflammatory pathologies suggest that dysregulation of NMD could contribute to a wider spectrum of diseases than previously appreciated ^41^.

Notably, the link between NMD disruption and innate immune activation may not be unique to the liver. For example, loss of the core NMD factor UPF2 in the mouse forebrain has previously been shown to trigger inflammation-associated phenotypes accompanied by innate and adaptive immune cell infiltration ^42^. Although the underlying mechanisms were not resolved in that study, these findings retrospectively support a broader connection between NMD integrity and immune homeostasis across tissues.

The strong immune activation induced by SMG6 loss also raises considerations regarding inflammatory toxicity. In *Smg6^mut^* livers, pronounced immune infiltration emerges at later stages, indicating that unchecked NMD inhibition can provoke tissue-specific inflammation. In an initial cohort, three *Smg6^mut^* mice died shortly after repeated blood sampling during weeks 7-8. Although no tumours were present at this early stage, the animals appeared unusually sensitive to procedural stress. Given the intense hepatic inflammation triggered by SMG6 loss, we consider it likely that acute cytokine-driven stress contributed to this vulnerability. Importantly, once blood sampling was discontinued, no further unexpected deaths occurred. These observations underscore the need to carefully evaluate inflammatory and toxicological consequences when considering pharmacological modulation of SMG6’s nuclease activity.

From a translational perspective, while SMG6 loss triggers dramatic immune activation in mice, our analysis of TCGA datasets indicates that *SMG6* mutations are rare in human cancers and generally appear as dispersed passenger events rather than PIN-domain-targeting alterations (**Supplementary Fig. 10a-b**). Analysis across different cancer types further shows no evidence of positive selection for *SMG6* mutations (**Supplementary Fig. 10c**). Because negative selection is statistically difficult to detect in tumour cohorts, the absence of enrichment cannot be interpreted directly; however, the lack of recurrent or domain-specific mutations is consistent with *SMG6* being required for tumour cell fitness. This interpretation is supported by large-scale CRISPR dependency screens, which show that complete *SMG6* loss is deleterious across a wide range of human cancer cell lines (**Supplementary Fig. 10d**).

A central limitation of the L-dKO genetic model is that SMG6 inactivation prevents tumour initiation entirely, precluding evaluation of SMG6 loss in established tumours. To address this and assess generalisability, we employed an orthotopic transplantation model. *Smg6-*deficient Hepa1-6 cells engrafted efficiently and initially grew at rates similar to wild-type cells, but subsequently exhibited significantly slower progression. This demonstrates that the tumour-restraining effects of SMG6 loss extend beyond tumour initiation and across distinct experimental contexts.

Finally, while only a limited number of non-canonical immunopeptides were identified in this genetically defined *Pten/Tsc1-*driven model, genetically heterogenous human tumours would be expected to harbour additional splice-altering or frameshift mutations. Such complexity would likely expand the repertoire of NMD-regulated neoantigens and further amplify adaptive immune responses triggered upon NMD inhibition.

Taken together, our data establish that loss of SMG6’s nuclease activity unmasks endogenous RNA species that activate MDA5-dependent type I IFN signalling and promote the presentation of non-canonical immunopeptides, creating a potent innate and adaptive immune environment that suppresses tumour growth. Although SMG6 inactivation prevents tumour initiation in the genetic model used here, our orthotopic transplantation experiment indicates that the immune-activating consequences of SMG6 targeting extend beyond tumour initiation. Future work across additional, genetically diverse HCC models will be important to determine the breadth of NMD-regulated neoantigens and to define how inflammatory toxicity can be controlled. Nonetheless, the mechanistic principles uncovered here provide a foundation for exploring selective, temporal, or partial inhibition of SMG6’s endonuclease activity as a means to increase tumour immune visibility in a clinically tractable manner.

## Methods

### Animals

The orthotopic tumour model was conducted under license AVD30100202519090 approved by the Dutch government and approved by the Dutch Animal Welfare Body of the Netherlands Cancer Institute (NKI). All other animal experiments were conducted in accordance with the cantonal guidelines of the Canton of Vaud, Switzerland, license VD3739. Healthy adult male mice of age 8 to 10 weeks were used. Animals with *Alb^CreERT2^*; *Smg6^wt^* and *Alb^CreERT2^*; *Smg6^flox^*alleles were maintained on a C57BL/6J background. Mice with *Alb^CreERT2^*; *Pten^flox^*; *Smg6^wt^*; *Tsc1^flox^* and *Alb^CreERT2^*; *Pten^flox^*; *Smg6^flox^*; *Tsc1^flox^* alleles were maintained on a hybrid background (C57BL/6J, 129SvJ, and BALB/2c). The alleles *Alb^CreERT2^*, *Pten^flox^*, *Smg6^flox^*, and *Tsc1^flox^* have been previously described ^20, 43–45^.

### Induction of hepatocyte-specific *Pten* & *Tsc1* double knockout and *Smg6* mutation

8- to 10-week-old male L-dKO *Smg6^wt^* and *Smg6^flox^*mice, carrying the liver-specific *Albumin*-driven *CreERT2*, and their control littermates received four intraperitoneal injections of tamoxifen, dissolved in corn oil at a dosage of 75 mg/kg of body weight, in the consecutive days. The early-stage mice were euthanised 3 weeks after tamoxifen injections. In the late-stage experiment, L-dKO *Smg6^wt^* mice were euthanised based on the progression of HCC (latest at week 17). Animals in the other groups (Ctrl *Smg6^wt^*, Ctrl *Smg6^mut^* and L-dKO *Smg6^mut^*) were euthanised 21 weeks after tamoxifen injections.

### Magnetic resonance imaging (MRI)

Mice were scanned weekly (starting from week 12), to monitor HCC evolution and progression, using a 3 Tesla small animal MRI scanner (Bruker BioSpin MRI, Ettlingen, Germany) with a 40-mm volume coil as a transmitter. Animals were anaesthetised using isoflurane (Attane) at 3-4% for 2 min. When fully sedated, animals were transferred into the MRI machine. The imaging procedure was performed under 1.5% isoflurane in oxygen. Data was acquired via Paravision 360 v2.0 software.

### Histopathology and immunohistochemistry analyses

Livers were fixed in 4% buffered formalin for 24 h and embedded in paraffin. For histology 3-5 µm-thick sections were cut and stained with Haematoxylin and Eosin (H&E) and Masson’s Trichrome (MTC). Immunohistochemistry was performed on an automated immunostainer (Ventana Medical Systems, Inc.) according to the company’s protocols for open procedures with slight modifications. All slides were stained with the antibodies CD4 (SP35; Zytomed, Berlin, Germany), CD8 (C8/144B; Dako, Glostrup, Denmark), PTEN (Cell Signaling Technology, 9188), Cleaved-Caspase3 (Cell Signaling Technology, 9661), and MPO (anti-MPO Ab-1, Lab Vision UK, Ltd., Newmarket, Suffolk, UK). Appropriate positive and negative controls were used to confirm the adequacy of the staining. The histologic samples were analysed by an experienced pathologist (L. Quintanilla-Martinez). All samples were scanned with the Ventana DP200 (Roche, Basel, Switzerland) and processed with the Image Viewer MFC Application. Final image preparation was performed with Adobe Photoshop 2024.

### Intrahepatic leukocyte isolation and flow cytometry staining

Leukocytes in liver were isolated as described previously ^46^. Mice were euthanised and subsequently perfused with 10 mL 1x PBS via the inferior vena cava. 1 g of liver tissue was excised from each mouse and placed into a tube containing 10 mL RPMI. The tissues were then crushed using 70 µm cell strainers and a plunger, ensuring all pieces were passed through the strainer. The resulting cell suspensions were centrifuged at 1600 × *g* for 5 min. The cell pellets were resuspended in a solution containing 2 mg collagenase, 25 µL DNase and 10 mL RPMI. The mixture was incubated at 37 °C for 40 min, constantly stirring. After incubation, 25 ml RPMI was added to dilute the mixture. The samples were then centrifuged at 300 × *g* for 3 min to precipitate the connective tissue. The supernatants were transferred to new tubes and centrifuged at 1600 × *g* for 5 min. The cell pellets were then resuspended in a Percoll density gradient solution (36% Percoll [Cytiva], 4% 10x PBS, 60% RPMI) and centrifuged at 2000 × *g* for 20 min without brake. The pellets were resuspended with 2 ml of Ammonium-Chloride-Potassium (ACK) lysis buffer for 2 min to lyse remaining red blood cells. Following that, 8 mL RPMI was added to the tubes, and the samples were centrifuged at 1600 × *g* for 5 min. The purified leukocyte pellets were resuspended in 1 mL HBSS. The cells were blocked with anti-CD16/32 and then washed with flow cytometry buffer (1x PBS, 2% FCS, 2 mM EDTA). Next, the cells were incubated with following primary antibodies for 25 min at 4 °C in the dark: CD19-BV605 (BD biosciences, 563148), TCRb-PE (BD biosciences, 12-5961-81), CD4-biotin (homemade), CD8a (Biolegend, 100750), CD44-FITC (homemade), CD11b-FITC (Biolegend, 101206), CD45-PE-Cy7 (Biolegend, 10114), Ly6C-BV785 (Biolegend, 128041), Ly6G-APC-Cy7 (Biolegend, 127624), CD19-PE (Immunotools, 22190194), TCRb-PE (BD biosciences, 12-5961-81), MHCII-BV711 (Biolegend, 107643), CD11c-biotin (homemade), NK1.1 (BV605, 108753), Streptavidin (APC, 405207) and BV510-Zombie aqua (Biolegend, 423102). Following washes with flow cytometry buffer, the cells were incubated with secondary antibodies for 20 min at 4 °C in the dark. The cells were fixed with BD fixation kit (554714).

### Serum parameters

Following the euthanasia of the animals, blood was quickly collected from the decapitated body, and then transferred into a lithium-heparin (LiHep) added tube (BD microtainer) to obtain serum. After 30 min to 1 h incubation at room temperature (RT), samples were centrifuged (2000 × *g*, 10 min, RT) to separate serum. Following that, 100 µL of serum was loaded to a Cobas c111 analyzer to measure the levels of following enzymes: ALT, AST and LDH.

### *Smg6* cDNA genotyping

Total RNA was prepared as described previously ^47^. For each RNA sample, the following reaction was assembled: 4.5 µg RNA, 1.5 µL random hexamer (100 µg/mL), 1.5 µL 10 mM dNTPs, RNase-free water to 18 µL. Samples were heated for 5 min at 65 °C, and then immediately put on ice. A master mix containing the following was added to each tube: 6 µL 5x First-strand buffer, 3 µL 0.1 M DTT, 0.375 µL RNase inhibitor, 1.5 µL RNase-free water. After a short incubation (2 min, at RT), 1.125 µL Superscript II reverse transcriptase enzyme was added to each reaction. Samples were incubated 10 min at 25 °C, 50 min at 42 °C, 15 min at 70 °C, respectively. *Smg6* genotyping polymerase chain reaction (PCR) was performed as described previously ^20^. The primer sequences – encompassing both mutant sites – are as follows: forward: CCA GAT ACC AAC GGC TTC AT; reverse: TCT GGG TGG ATT GGT AGC TC. Reactions were run on a 0.8% agarose gel, at 100 V for 100 min. Bands corresponding to ∼620 nt were excised, and then purified using QIAquick Gel Extraction Kit (QIAGEN, 28704). DNAs were Sanger-sequenced by Microsynth.

### Quantitative Reverse Transcriptase PCR

Total RNA was prepared, and cDNA was synthesised as mentioned above. 10-times diluted cDNA templates and a master mix containing SYBR green master mix (2x), forward and reverse primers (3.0 µM) were mixed, and then a 384-well plate was prepared using the Tecan Freedom EVO software. RT-PCR was performed using QuantStudio6 real-time systems. Relative mRNA expression levels were obtained by normalizing each CT value (mean of three technical replicates) to *Rpl13* and *Rpl27* expressions. The qPCR primer sequences can be found in **Supplementary Table 2**.

### Transcriptomics analysis

Total RNA was prepared as mentioned above. RNA-seq libraries were prepared from individual mouse livers according to the standard protocols. Single-end sequencing (150 bases) was performed on NovaSeq6000 (Illumina) with a sequencing depth of 400 million cDNA-mapping reads in total. Sequencing reads were trimmed for the first two low quality (nucleotides) nt. Trimmed reads were then mapped using STAR ^48^ (v2.7.11b; --twopassMode Basic --seedSearchStartLmax 50 --outSAMmultNmax 1) in a sequential manner: mouse rRNA (Ensembl version 39.111), human rRNA (Ensembl version 38.111), mouse tRNA, and mouse genome. Read counts per genes were obtained using summariseOverlaps() of the R (v4.4.0) library GenomicAlignments (mode = ’Union’) and low read counts were excluded (≤5 counts in less than three samples). Differential gene expression analysis was performed using DEseq2 ^49^.

### Ribosome profiling

The libraries for ribosome profiling were generated in principle as described ^50^ with small adaptations. Frozen liver tissues (200 mg) from individual mice were homogenised with 5 strokes using a Teflon homogeniser in 3 volumes of polysome buffer (150 mM NaCl, 20 mM Tris-HCl [pH 7.4], 5 mM DTT, complete EDTA-free protease inhibitors [Roche] and 40 U/mL RNasin plus [Promega]) supplemented with 1% Triton X-100 and 0.5% Na deoxycholate. Liver lysates were incubated on ice for 10 min and cleared by centrifugation at 10000 × *g*, 4 °C for 10 min. Supernatants were flash-frozen and stored in liquid nitrogen. For each sample, 15 OD_260_ of liver lysate was digested with RNase I (Ambion) for 45 min at RT under constant shaking. RNase I-digested RNA was separated on 15% urea-polyacrylamide gels to excise monosome (∼30 nt) footprints. From the gel slices, RNA was extracted overnight at 4 °C on a rotating wheel before precipitation in isopropanol for 1 h, at -20°C. RNA 3’-end repair was performed with 2 U/µL of T4 PNK (Lucigen) before 2 h of adapter ligation at 25 °C using T4 RNA Ligase 1 (NEB) and T4 RNA Ligase 2 Deletion Mutant (Lucigen) and 1 µL of a 20 µM 5’-adenylated DNA adapter. Adapter removal was performed by treating individual libraries with 5’ deadenylase (NEB) and RecJf exonuclease (NEB) for 1 h at 30°C and 1 h at 37°C. Samples were purified and pooled using Zymo Clean & Concentrator columns. Ribosomal RNA was depleted according to siTools Biotech rRNA depletion kit specifications with a custom-made riboPOOL. The clean-up step was performed using Zymo Clean & Concentrator columns. Further library preparation steps were performed as described. The amplification of libraries was performed using i5 and i7 [NexteraD502 or NexteraD503] primers. Libraries were sequenced on a NovaSeq6000 (Illumina).

### Ribosome profiling data analysis

Sequencing reads were cleaned and trimmed. Briefly, adapter sequences were removed using cutadapt (v3.5) ^51^ with the following parameters --match-read-wildcards --overlap 8 --discard-untrimmed --minimum-length 30. Reads were filtered by quality using FASTX Toolkit (v0.0.14; -Q33 -q 30 -p 90) and by size using an in-house Python script (length 26-35 nt). Cleaned reads were then mapped using STAR (--twopassMode Basic –seedSearchStartLmax 28 – outSAMmultNmax 1) ^48^ in a sequential manner: mouse rRNA (Ensembl version 39.111), human rRNA (Ensembl version 38.111), mouse tRNA, and mouse genome. STAR was used for the mapping to splice junctions, as neoantigens can be produced from both exons and introns. The A-site position of mapped reads to the mouse genome was retrieved using the R (v4.4.0) package ribosomeProfilingQC (v1.16.0; estimatePsite() and getPsiteCoordinate(), and addition of 3 nt to recover the A-site position). The A-sites were assigned to the counting bins and the A-site count per bin was retrieved. Counting bins were defined using an in-house R script (main functions: makeTxDbFromGFF() from txdbmaker [v1.1.1]; exonicParts() from GenomicFeatures [v1.57.0]; and reduce(), gaps(), and findOverlaps() from IRanges [v2.39.2]) and A-site counts were obtained using summariseOverlaps() of GenomicAlignments (v1.41.0; mode = ‘Union’) ^52^. The low A-site counts (≤5 counts in less than three samples) were excluded and the differential analysis was performed using DESeq2 (v.1.45.3). To identify putative neoantigens, the median A-site counts were calculated for each condition and the bins with at least one A-site count in L-dKO *Smg6^mut^* were retained. The nucleotide sequences of each bin were extracted, translated in the three reading frames, and the resulting amino acid sequences were then split by stop codon, and retained if the peptide sequence lengths were ≥ 8 amino acids.

### MHC-I antibody production

MHC-I monoclonal antibodies were purified from the supernatant of HB-51 and HIB 34.1.2 hybridoma cells (ATCC), respectively, with Protein A-Sepharose 4B beads (Invitrogen). HB-51 has specificity for H2-Db and H2-Kb, HIB 34.1.2 has specificity for H2-Dd, H2-Kd, and H2-Ld. Each antibody type was cross-linked separately to Protein A-Sepharose 4B beads at concentrations of 5 mg of antibodies per 1 mL of bead volume. To this end, the antibodies were incubated with Protein A-Sepharose 4B beads for 1 hour, at RT. Antibodies and beads were cross-linked with dimethyl pimelimidate dihydrochloride (Sigma-Aldrich) in 0.2 M sodium borate buffer pH 9.0 (Sigma-Aldrich) to achieve a final concentration of 20 mM for 30 min. The reaction was quenched via incubation with 0.2 M ethanolamine pH 8.0 (Sigma-Aldrich) for 2 hours. Cross-linked antibodies were stored at 4 °C.

### Purification of MHC-I-eluted peptides

200 mg of liver tissue per biological replicate was lysed in 2 mL phosphate buffered saline with 0.5% sodium deoxycholate (Sigma-Aldrich), 0.2 mM iodoacetamide (Sigma-Aldrich), 1 mM EDTA, 1:200 Protease Inhibitor Cocktail (Sigma-Aldrich), 1 mM phenylmethylsulfonylfluoride (Roche), and 1% octyl-beta-D glucopyranoside (Sigma-Aldrich) at 4 °C for 1 hour, after homogenization with an IKA T10 standard ULTRA TURRAX. Lysates were cleared by centrifugation with a table-top centrifuge (Eppendorf Centrifuge) at 4 °C, 20000 × *g* for 50 min. MHC-I molecules purification was done with HB51 antibodies for C57BL/6 and a 1:1 mix of HB51, HiB antibodies for C57BL/6; BALB/2c; 129_SvJae. Immunoprecipitations in 96-well plates followed the protocol described before ^53^. Shortly, after plate conditioning, cross-linked beads and then lysates were added to single wells, followed by washes at different sodium chloride concentrations and an acidic elution step. Peptides were then desalted and purified on C18-columns. Recovered peptides were dried using vacuum centrifugation (Thermo Fisher Scientific) and stored at −80 °C.

### Mass-spectrometry acquisition

For Mass spectrometry (MS) analysis, MHC-I peptide samples were resuspended in 8 µL of 2% ACN and 0.1% formic acid (FA). 3 µL of peptide resuspension was injected into the LC-MS setup per measurement, counting two in DDA and one in DIA. The LC-MS setup consists of an Easy-nLC 1200 (Thermo Fisher Scientific) connected to a Q Exactive HF-X (Thermo Fisher Scientific). Chromatographic separation was done with a 450 mm homemade column of 75 µm inner diameter packed with ReproSil Pur C18-AQ 1.9 µm resin (Dr. Maisch GmbH). For DDA measurements, the chromatographic separation was performed over 2 hours using a gradient of H_2_O:FA, 99.9%:0.1% (solvent A) and CH_3_CN:FA 80%:0.1% (solvent B). The gradient consisted of sequential, linear steps: 0 to 110 min (2-25% B), 110 to 114 min (25-35% B) , 114 to 115 min (35-100% B), and 115 to 120 min (constant at 100% B) at a flow rate of 250 nl/min. The mass spectrometer had the following parameters set: full-scan MS spectra were acquired from m/z = 300-1,650 at a resolution of 60,000 (m/z = 200) with maximum injection times of 80 ms. Auto gain control (AGC) target value was tuned to 3 x 10^6^ ions. We acquired MS/MS spectra at a resolution of 30,000 (m/z = 200), a top 20 method and isolation windows of 1.2 m/z, as well as a collision energy of 27 (HCD). All MS/MS scans accumulated ions to an AGC target value of 2 x 10^5^ at a maximum injection time of 120 ms. Precursors of charge 4 and above were not selected for fragmentation. The dynamic exclusion was set to 20s. For DIA, the chromatographic separation was performed as in DDA measurements. The DIA acquisition cycles included one full-scan MS spectra, acquired from m/z = 300-1,650 (Resolution = 120,000, ion accumulation time = 60 ms), and 22 MS/MS scans. For all MS/MS scans the AGC target value was set to 3 x 10^6^ ion, mass resolution at 30,000 and a stepped normalised collision energy (25.5, 27, 30) was applied. The maximum ion accumulation was set to auto and their first mass was fixed to 200 m/z. Consecutive MS/MS scans overlapped in their m/z windows by 1 m/z.

### Immunopeptidomics data processing

All DDA files were used to generate a library with a database search using FragPipe (v22.0), including MSFragger (v4.1) ^54^. The FASTA file was compiled as described in ‘Ribosome profiling data analysis’. For search space generation, unspecific digestion with peptide lengths of 8 to 14 amino acids was used. Variable modifications included methionine oxidation, N-terminal acetylation, and cysteine carbamidomethylation. No fixed modifications were applied. Peptide, ion, and psm FDRs were set to 0.03 while applying a group-specific FDR as indicated in the FASTA header with PE = X, X being the respective protein group. No protein FDR was applied. All other settings were set to default. DIA files were searched with the library described above using DIA-NN (v1.8) with a precursor q-value cut-off of 0.01. Peptide binding affinities were predicted with MixMHC ^55^. Peptides were considered as binders when they had a %-rank below 2 for at least one HLA allele present in the sample haplotypes. Data analyses were done with Julia and R, visualizations with R.

### Protein extraction and western blot

Total proteins from mouse liver samples were isolated according to the NUN (NaCl, Urea, Nonidet P-40) procedure, as ^20^: Livers were homogenised in two tissue volumes of 10 mM Hepes (pH 7.6), 100 mM KCl, 0.1 mM EDTA, 10% glycerol, 0.15 mM spermine, and 0.5 mM spermidine for 15 s using a Teflon homogeniser. Four tissue volumes of 2x NUN buffer (2 M urea, 2% NP-40, 0.6 M NaCl, 50 mM Hepes [pH 7.6], 2 mM dithiothreitol, and 0.1 mM phenylmethylsulfonyl fluoride; supplemented with complete protease inhibitor tablets, Roche) were added dropwise on a vortex with low speed. Lysates were incubated 1 h on ice and cleared through centrifugation at 10,000 rpm, 4°C, for 20 min. Aliquots of the lysates (40 µg of protein loaded per lane) were separated by SDS-polyacrylamide gel electrophoresis and transferred to PVDF membrane using an iBlot 2 gel transfer device. The membrane was blocked using 5% milk in Tris-buffered saline with 0.1% Tween 20 (TBST), 1 hour, at RT, followed by an incubation overnight at 4 °C with appropriate dilutions of primary antibodies, including anti-Cleaved CASPASE3 (Cell Signaling Technology, 9661), anti-MDA5 (Cell Signaling Technology, 5321), anti-PTEN (Cell Signaling Technology, 9188), anti-Hamartin/TSC1 (Cell Signaling Technology, 6935), anti-GAPDH (Cell Signaling Technology, 2118), and anti-VINCULIN (Abcam, ab129002). Following TBST washing (5 min, three times), the membrane was incubated with the appropriate secondary antibody conjugated with horseradish peroxidase for 60 min at RT, followed by TBST washing (15 min, three times). Chemiluminescence signal was detected using Supersignal West Femto Maximum Sensitivity Substrate (Thermofisher Scientific, 34095), as described by the manufacturer. The quantification of signal was performed using ImageJ software.

### Deletion of *Ifih1* and *Smg6* via CRISPR

*Ifih* was deleted in *Smg6^flox/flox^*; *CreERT2* fibroblasts while *Smg6* was knocked out in Hepa1-6 cells. First, lentiCRISPRv2-Blast-mU6 empty backbone (Addgene #206806) ^56^ was digested with BsmBI (NEB) enzyme. The digestion products were run on an agarose gel (0.8%), and the band corresponding to the open vector (∼11 kb) was excised, followed by DNA purification using QIAquick Gel Extraction Kit. Following annealing reaction was assembled in a tube: 1 µL Forward primer (100 µM), 1 µL Reverse primer (100 µM), 1 µL 10x T4 ligation buffer (NEB, B0202S), 0.5 µL T4 PNK (NEB, B0201S), and 6.5 µL H_2_O. Primers were annealed with the following protocol: incubate 30 min at 37 °C, and 5 min at 95°C, respectively, then cool to 25 °C at a rate of 5°C/min. Annealed primers were ligated into the open vector using the following mixture: 50 ng open vector, 2 µL 10x T4 ligation buffer, 1 µL T4 DNA ligase (NEB, M0202S), 1 µL annealed primers (1:100 diluted), and nuclease-free H_2_O to 20 µL. Ligation mix was incubated at RT for 1 h. 10 µL ligation mix was added to 100 µL Stbl3 competent cells, and incubated on ice for 30 min, followed by a heat-shock for 30 sec at 42 °C. After a short incubation (5 min, on ice), 1 mL S.O.C. medium was added to the mixture, and the tube was incubated at 37 °C, for 1 h with shaking. 100 µL of the transformation mixture was plated on an agar plate containing Ampicillin and incubated overnight at 37 °C. Positive colonies were selected and cultured overnight with shaking at 37 °C, in LB medium containing Ampicillin, then purified using a Midiprep kit (QIAGEN, 12943). Cells were transfected with the plasmid using Lipofectamine 3000 (Thermo Fisher Scientific, L3000001), according to the manufacturer’s protocol, for 24 h, then were selected with Blasticidin (5 µg/mL) for 96 hours. Knockout cells were obtained through single cell cloning. *Ifih1* sgRNA primer sequences: forward, CAC CGC GTA GAC GAC ATA TTA CCA G; reverse, AAA CCT GGT AAT ATG TCG TCT ACG C. *Smg6* sgRNA primer sequences: forward, CGT CAG GCC CAA GCA TCT TA; reverse, AGG GAG AGG GCT ACA CAC AT.

### Induction of Cre-mediated recombination in fibroblasts

Two different methods were used to induce Cre-dependent recombination. In the first series of experiments, we used 4-OHT to activate retrovirally expressed CreERT2 (i.e. in **Fig. 4c-e**). In clonal cell lines derived later, where CreERT2 transgene expression can be subject to clonal differences, the TAT-CRE system provided more reliable delivery (i.e. in **Fig. 5b-f**).

i. Direct delivery of recombinant CRE protein (TAT-CRE, kind gift from I. Lopez-Mejia)*: Smg6^w/wt^* and *Smg6^flox/flox^*(-/+ *Ifih1^ko/ko^*) fibroblasts were seeded in 6-well plates. The standard culture medium (Thermo Fisher Scientific, 41965062) was replaced with Opti-MeM (Thermo Fisher Scientific, 31985062). Cells were subsequently treated with 2.5 µM TAT-CRE or vehicle for 8 h. After treatment, OptiMeM was replaced with the standard culture medium.
ii. Recombination via 4-hydroxytamoxifen (4-OHT): *Smg6^flox/flox^*; *CreERT2* cells were treated with 2 µM 4-OHT or DMSO for 3 days (4-OHT containing media was refreshed every 24 h). In both methods, cells were maintained in culture for at least one-week post-treatment to ensure efficient recombination. Cells were subsequently collected for RNA extraction (as explained above).

### Inhibition of NMD in *Smg6^flox^* fibroblasts via SMG1 inhibitor

*Smg6^flox^*primary fibroblasts were treated with SMG1 inhibitor (hSMG-1 inhibitor 11e, ProbeChem PC-35788, dissolved in DMSO) at a concentration of 0.6 µM, during 24 h.

Following the treatment, the cells were collected and then RNA was isolated as explained above. NMD inhibition was confirmed via increased transcript abundance of the NMD-target variant of *Hnrnpl* using qPCR.

### J2 immunofluorescence

Immunofluorescence of dsRNA was performed as described ^57^. First, *Smg6^flox/flox^* cells were seeded in a 6-well plate, then treated with 2.5 µM TAT-CRE or vehicle as above. One week after the treatment, cells were transferred to a new 6-well plate containing coverslips, at a confluency of ca. 20%. Next day, the culture medium was removed, and cells were fixed using 4% paraformaldehyde (PFA) for 15 min at RT, then washed three times (5 min each time), with 1x PBS. Fixed cells were permeabilised with 500 µL PBSTX, followed by a 10 min incubation at RT. PBSTX was removed, 500 µL of IFA blocking buffer was added into each well, and incubated for 1 h, at RT with rocking. After blocking, cells were incubated with 500 µL primary antibody solution (anti-dsRNA monoclonal antibody J2: Jena Bioscience, RNT-SCI-10010200) in IFA blocking buffer (1:1000), overnight, at 4°C with rocking. Primary antibody solution was removed, and cells were washed with 500 µL PBSTX (three times, for 5 min each). Cells were incubated with the secondary antibody solution in PBSTX (1:300, Goat anti-mouse IgG Alexa Fluor 488 for detection of J2, followed by washes with 500 µL PBSTX 3 times, for 5 min each). To prepare the slides for imaging, coverslips were removed from 6-well plate, and cleaned with 70% ethanol, then placed face down onto the glasses, sealed with a nail polish. Images were captured using a fluorescence microscope. The quantification of J2 intensity was performed on ImageJ software.

### Enzyme-linked immunosorbent assay (ELISA)

CXCL10 ELISA (R&D, DY466) was performed using R&D DuoSet^TM^ ELISA development system, according to manufacturer’s protocol. 96-well ELISA plate (R&D, DY990) was coated with 100 µL of capture antibody solution and incubated overnight at room temperature. Following washes using wash buffer (R&D, WA126) three times, plate was blocked with 300 µL block buffer (R&D, DY995) for 1 hour at room temperature. Washes were repeated as above. 100 µL of diluted samples and standards were added to the plate and then incubated for 2 hours at room temperature. Washes were repeated as above. 100 µL of detection antibody solution was added to the plate and then incubated for 2 hours at room temperature. Washes were repeated as above. 100 µL of Streptavidin-HRP B solution was added to the plate and then incubated for 20 minutes at room temperature. Washes were repeated as above. 100 µL of substrate solution (R&D, DY999B) was added to the plate and then incubated for 20 minutes at room temperature. The reaction was stopped via stopping solution (R&D, DY994). Protein levels were detected using a microplate reader at 450 nm optical density.

IFN-α and IFN-β levels were determined using Lumikine^TM^ Xpress ELISA system (mIFN-α, luex-mifnav2; m IFN-β, luex-mifnbv3). 96-well white flat bottom plate (provided with the kit) was coated with 50 µL of capture antibody solution overnight at room temperature. After removing the antibody solution, the plate was blocked with 200 µL of blocking buffer (1x PBS containing 3% BSA and 0.05% Tween 20) during 2 h at 37°C. Then, blocking buffer was removed and 50 µL of samples (or standards) and 50 µL of Lucia-conjugated detection antibody solution were added to the plate, respectively. The plate was incubated during 2 h at 37°C. Following three washes with the wash buffer (1x PBS containing 0.05% Tween 20), 50 µL of QUANTI-Luc^TM^ 4 Reagent working solution was added to the plate. Bioluminescent signal was immediately measured in a plate reader (Tecan Safire 2) at 0.1 second reading time.

### Ex vivo peptide stimulation of leukocytes

Intrahepatic leukocytes isolated from mice as explained above. Spleens were mechanically disrupted and filtered through a 70 µm cell strain to obtain single-cell suspensions. Both intrahepatic leukocytes and spleens were washed with PBS. 2 million cells per mouse per condition were seeded in round-bottom 96-well plates in T-cell medium (RPMI supplemented with 25 µM 2-mercaptoethanol, 1 mM sodium pyruvate, MEM non-essential amino acids (Gibco^TM^, 11140050), 10% FBS, 1% Penicillin/Streptomycin, and L-glutamine. In the first assay, a pool of non-canonical, canonical, and common peptides was added to the appropriate wells at a final concentration of 2 µg/mL per peptide (Fig. 3f-h; Supplementary Fig. 4b-c). In the second assay, cells were incubated with four non-canonical peptides either separately or in a pool at 2 µg/mL concentration per peptide (Supplementary Fig. 5a-d). CD28 agonist antibody (1 µg/mL; BioLegend, 102115) and Brefeldin A were added to all wells. Cells were incubated at 37°C for 6 hours. After incubation, cells were collected and stained with Zombie UV^TM^ Fixable Viability Kit (1:500; BioLegend, 423107) to stain dead cells. Then, extracellular and intracellular staining was performed using BD Cytofix/Cytoperm^TM^ Fixation/Permeabilization Kit (Cat# 554714) according to the manufacturer’s instructions. Data were acquired on a BD LSR Fortessa flow cytometer and analyzed using FlowJo software. Antibodies used were: CD3-PE (BioLegend, 100206), CD45-Pacific Blue (BioLegend, 103126), CD8a-Brilliant Violet 510 (BioLegend, 100714), and IFNγ-APC (BioLegend, 505810).

### RNA structure prediction

RNA structure prediction was performed via RNAfold web server using default parameters (http://rna.tbi.univie.ac.at/cgi-bin/RNAWebSuite/RNAfold.cgi).

### Repetitive element analysis

Repetitive elements within individual transcripts were detected using RepeatMasker ^58^ web server (https://www.repeatmasker.org/cgi-bin/WEBRepeatMasker).

### Cell cycle analysis by propidium iodide (PI) staining

Hepa1-6 cells were seeded at 1.5 x 10^5^ cells per well (6-well plates) and cultured for 48 h to reach ∼70% confluency. Then, cells were harvested by trypsinization, transferred into FACS tubes, pelleted (1200 rpm, 3 min), washed twice in 1x PBS, and permeabilised using 1x saponin-based permeabiliation solution (Click-iT kit EdU, C10424, Invitrogen) at room temperature in the dark. A final wash in the same buffer was performed before staining. For DNA content analysis, cells were incubated for 30 min at room temperature in the dark in 1x saponin buffer and analysed by flow cytometry. Cell cycle distribution was quantified using FlowJo with the Watson Pragmatic model.

### Orthotopic tumour implantation experiment

Animals were housed in individually ventilated cages with access to food and water *ad libitum*. 24 wild-type immunocompetent male C57BL/6 mice were inoculated orthotopically in the liver, with ultrasound guided injections (UGI) on the Visual Sonics Vevo F2, with 50 µl 1 x 10^6^ Hepa1-6 (n = 12 for *Smg6^wt/wt^*; n = 12 for *Smg6^ko/ko^*) tumour cells (resuspended in Matrigel (Corning)). Mice were anaesthetised with 2-4% isoflurane and body temperature was monitored. Starting from the first week, tumour development was monitored by MRI twice per week. MRI was performed in ParaVision 7.0.0 on a 7T Bruker BioSpec 70/20 USR with a 1H transmit-receive volume coil. T_2_-weighted images were acquired under 1-2% isoflurane in air and oxygen flow using a respiratory-gated sequence with TR/TE = 2,500/25 ms, 32 x 24-mm field of view (320 x 240 matrix, resolution of 0.1 mm), 30 x 0.7 mm axial slices and 4 averages. Mice were weighted daily. 4 animals from each group were sacrificed 3 weeks after injections for early-stage comparison and the remaining mice were sacrificed 5 weeks after injections. Tumour volumes were calculated via 30 abdominal MRI slices per animal for each timepoint using Bee DICOM viewer software. Following euthanasia of the animals, livers were perfused with 10 ml 1x PBS via inferior vena cava. Liver tumours were either stored at −80°C for RNA and protein extractions, or fixed with formalin, and embedded in paraffin for histological examination.

### TCGA analysis on *SMG6* mutation occurrence in different cancers

Mutation data for *SMG6* were obtained from the TCGA PanCancer Atlas cohort ^59^ via the cBioPortal for Cancer Genomics ^60^. Mutation frequency, variant type, and predicted oncogenicity were extracted and summarized across all tumour types. Protein level positional distributions of missense mutations were visualized to assess clustering or hotspot formation. To evaluate evidence of positive selection, dN/dS analysis ^61^ was performed using only tumour types with ≥10 *SMG6*-mutated samples. Additionally, CRISPR-based dependency scores for *SMG6* were retrieved from DepMap (DepMap, Broad, Public 25Q3) to assess the essentiality of the gene across cancer cell lines ^62^.

## Acknowledgements

We thank the Lausanne Genomic Technologies Facility for library preparation and sequencing, University of Lausanne Peptide & Tetramer Core Facility for the synthesis of synthetic peptides, F. Lammers for help with genotyping, C. Roger for histology analysis, G. Niederhäuser for serum measurements, Anne-Claire Godet at www.threoninedesign.fr for the graphical illustration in Figure 5g, I. Lopez-Mejia for providing TAT-CRE, J. Van Leeuwen for providing lentiCRISPRv2-Blast-mU6 plasmid, M. van de Ven, N. Proost, and M. Boeije from the Netherlands Cancer Institute for the orthotopic tumour implantation experiment, O. Mühlemann and E. Karousis for important discussions. D.G. acknowledges funding by Swiss National Science Foundation (SNSF) NCCR RNA & Disease Phase III funding (grant number 205601), SNSF project grant (212423) and by Novartis Foundation for Medical-Biological Research grant (21B066).

## Author Contributions

Conceptualization: E.S.A. and D.G.; methodology: E.S.A., V.R., I.E.S., M.J., Y.M., L.B., I.G.- M., and A.P.B.; investigation: E.S.A., V.R., I.E.S., M.J., Y.M., I.G.-M., L.Q.-M., and D.M.; visualization: E.S.A., V.R., I.E.S., M.J., Y.M., I.G.-M., and G.C.; supervision: D.G., L.Q.-M., M.N.H., J.R., T.V.P., S.A.L., and M.B.-S.; writing: E.S.A. and D.G.

## Data Availability

The transcriptomics and ribo-seq data discussed in this publication have been deposited in NCBI’s Gene Expression Omnibus ^63^ and are accessible through GEO Series accession numbers GSE278996 and GSE278997. The mass spectrometry proteomics data have been deposited to the ProteomeXchange Consortium via the PRIDE ^64^ partner repository with the dataset identifier PXD057903.

## Code Availability

All computational scripts are available on GitHub (https://github.com/NMD_Smg6/).

---

**Supplementary Figure 1.**
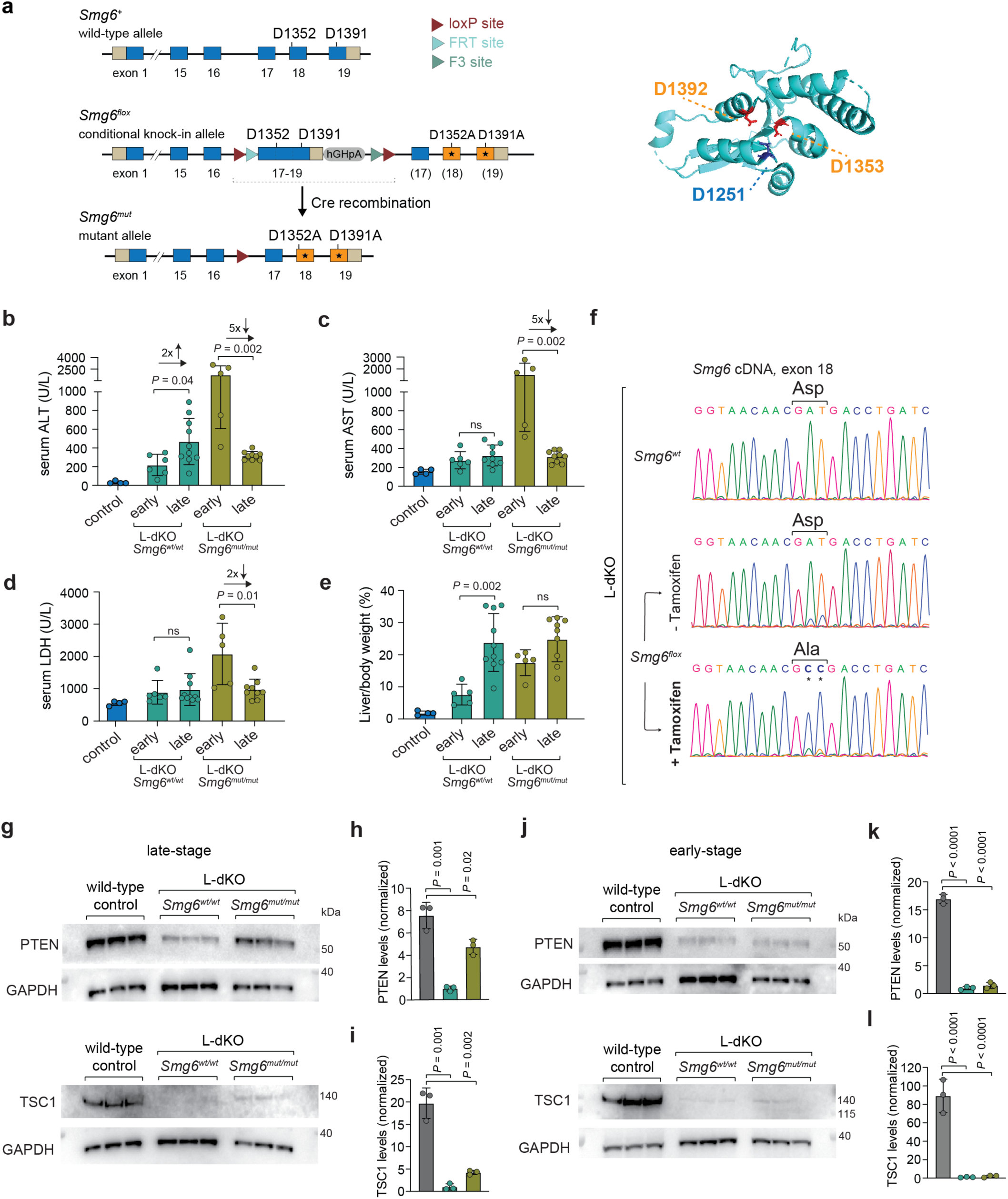
Regression of liver damage marker levels in *Smg6* mutant mice from early- to late-stage L-dKO. (a) Schematic of the genetic NMD-deficient mouse allele based on *Smg6*. *Smg6^flox^* expresses wild-type SMG6 protein encoded by the blue exons; after Cre-mediated recombination to *Smg6^mut^*, point-mutated exons 18 and 19 (orange) lead to the expression of mutant SMG6 (D1352A, 1391A). The right panel shows the structure of the human protein (PDB accession 2HWW) with the corresponding mutated aspartic acid residues (in orange) that are within the catalytic triade of the PIN nuclease domain. Please note that human Asp1392 corresponds to mouse Asp1391, and human Asp1353 to mouse Asp1352. Figure adapted from ^16^. (b-d) Comparison of the levels of indicated liver damage enzymes, detected in serum, between early- and late-stage mice. Arrows display the fold-change alteration in the level of liver damage markers from early- to late-stage, in the same group (n = 4 mice for control, n = 5 mice for early-stage groups; L-dKO *Smg6^wt^* late, n = 10 mice; L-dKO *Smg6^mut^* late, n = 9 mice). (e) Comparison of the ratio of liver-over-body weight between early- and late-stage mice, within the same group (n = 5 mice for early-stage groups; L-dKO *Smg6^wt^* late, n = 10 mice; L-dKO *Smg6^mut^* late, n = 9 mice). (f) Sanger sequencing chromatograms on cDNA, from representative late-stage livers, show the conversion of aspartic acid (Asp) encoding wild-type codon (GAT) to alanine (Ala) encoding mutant codon (GCC), at the position in exon 18 that codes for one of the Asp residues of the catalytic center of SMG6 that was targeted in the genetic model, through tamoxifen-induced CreERT2 activation in liver. (g) Western blot analysis of total liver proteins from late-stage mice for PTEN and TSC1. GAPDH served as loading control (n = 3 mice for each group). (h) Quantification of PTEN and GAPDH western blots from (g). (i) Quantification of TSC1 and GAPDH western blots from (g). (j) Western blot analysis of total liver proteins from early-stage mice for PTEN and TSC1. GAPDH served as loading control (n = 3 mice for each group). (k) Quantification of PTEN and GAPDH western blots from (j). (l) Quantification of TSC1 and GAPDH western blots from (j). For (b), (c), (d), (e), (h), (i), (k), and (l) data are plotted as means, and the error bars indicate SD. The *P*-values were calculated with two-tailed unpaired *t*-test.

**Supplementary Figure 2.**
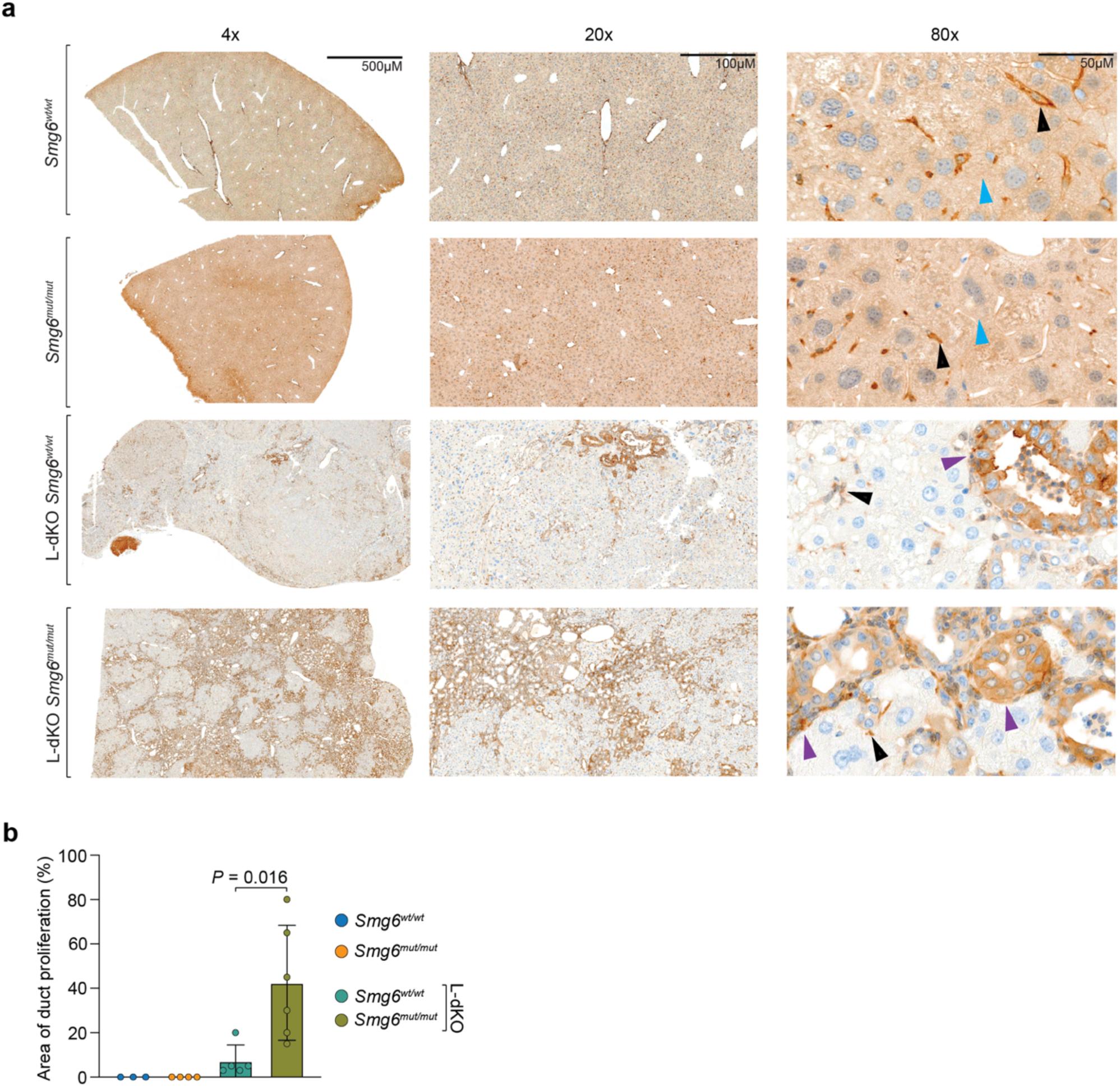
L-dKO leads to duct proliferation in liver that is more aggravated in *Smg6^mut^*and that contributes most PTEN signal. (a) Representative IHC images of livers from late-stage mice stained with anti-PTEN antibodies. Brown arrows indicate selected non-hepatocyte cells with strong PTEN staining. Purple arrowheads show L-dKO-specific duct proliferation, which is more pronounced in *Smg6^mut^*tissues. Of note, PTEN staining is homogenously present in non-L-dKO hepatocytes (upper two rows), yet specifically absent from L-dKO hepatocytes, as expected. (b) Quantification of area showing duct proliferation. For each sample, the area of duct proliferation was calculated in 15 randomly selected images per liver, at 20x magnification (*Smg6^wt^*, n = 3 mice; *Smg6^mut^*, n = 4 mice; L-dKO *Smg6^wt^*, n = 5 mice; L-dKO *Smg6^mut^*, n = 6 mice). Data are plotted as means, and the error bars indicate SD. *P*-value was calculated with two-tailed unpaired *t*-test.

**Supplementary Figure 3.**
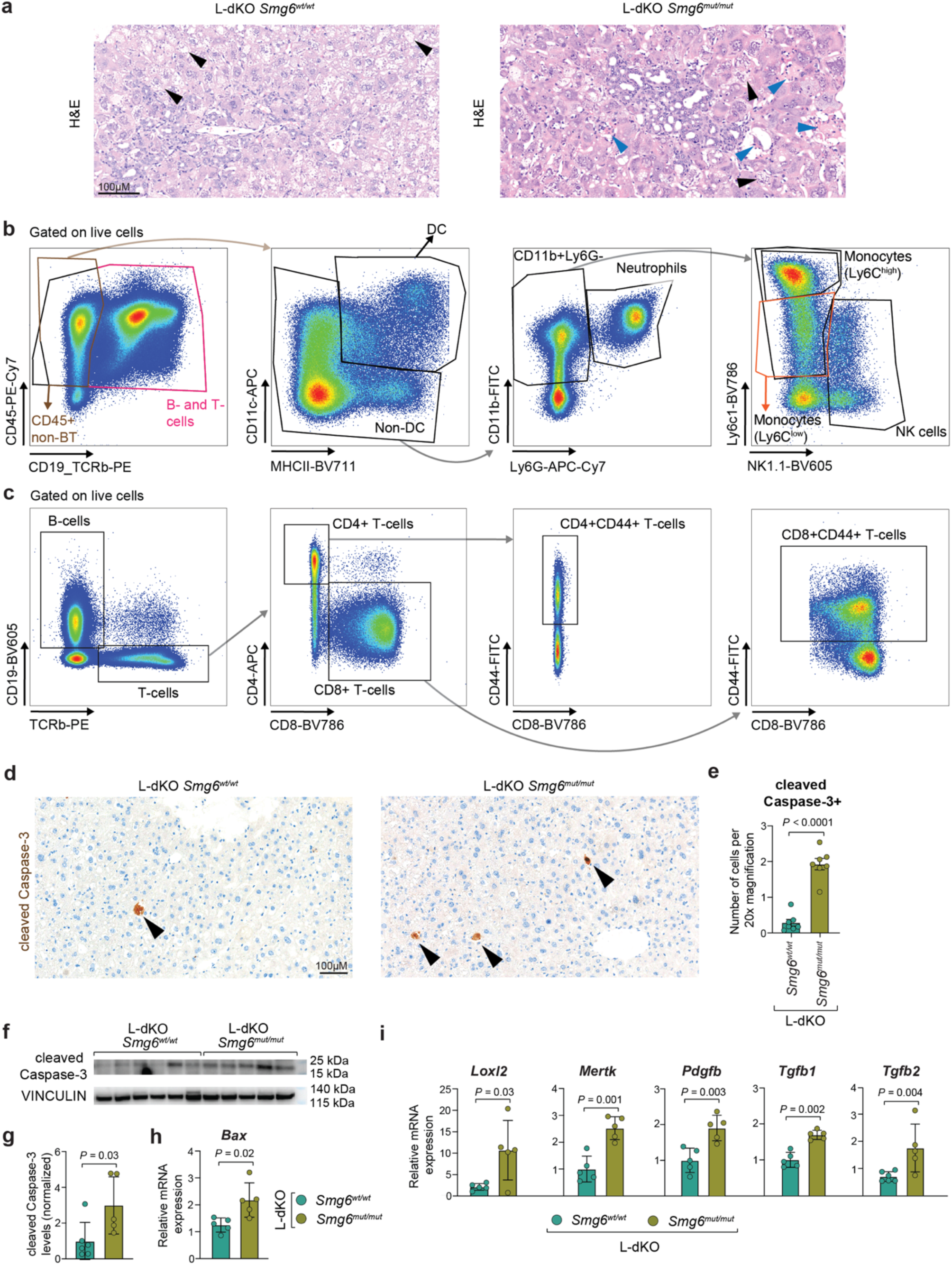
Cellular changes following *Smg6* mutation in liver. (a) Representative H&E images from early-stage L-dKO livers, at 20x magnification. Black arrowheads indicate hepatocyte ballooning in both livers, while blue arrowheads mark apoptotic hepatocytes in L-dKO *Smg6^mut^*tissue. (b) Flow cytometry gating strategy of intrahepatic myeloid cells. (c) Flow cytometry gating strategy of intrahepatic lymphocytes. (d) Representative IHC images of livers from early-stage mice, stained with antibodies directed against cleaved Caspase-3, at 20x magnification. Arrowheads show cleaved Caspase-3+ cells (brown). (e) Quantification of cleaved Caspase-3 IHC images as that shown in (d). For each sample, the number of cleaved Caspase-3+ cells was counted in 10 randomly selected images, at 20x magnification (mean values from 7 mice per group). (f) Western blot analysis of total proteins, for cleaved Caspase-3. VINCULIN served as loading control (L-dKO *Smg6^wt^*, n = 6 mice; L-dKO *Smg6^mut^*, n = 5 mice). (g) Quantification of western blot from (f). (h) Relative hepatic mRNA levels of *Bax*, from early-stage mice (n = 5 mice per group). (i) Relative hepatic mRNA levels of indicated pro-fibrotic genes, from early-stage mice (n = 4 mice per group). For (e), (g), (h), and (i) data are plotted as means, and the error bars indicate SEM. The *P*-values were calculated with two-tailed unpaired *t*-test.

**Supplementary Figure 4.**
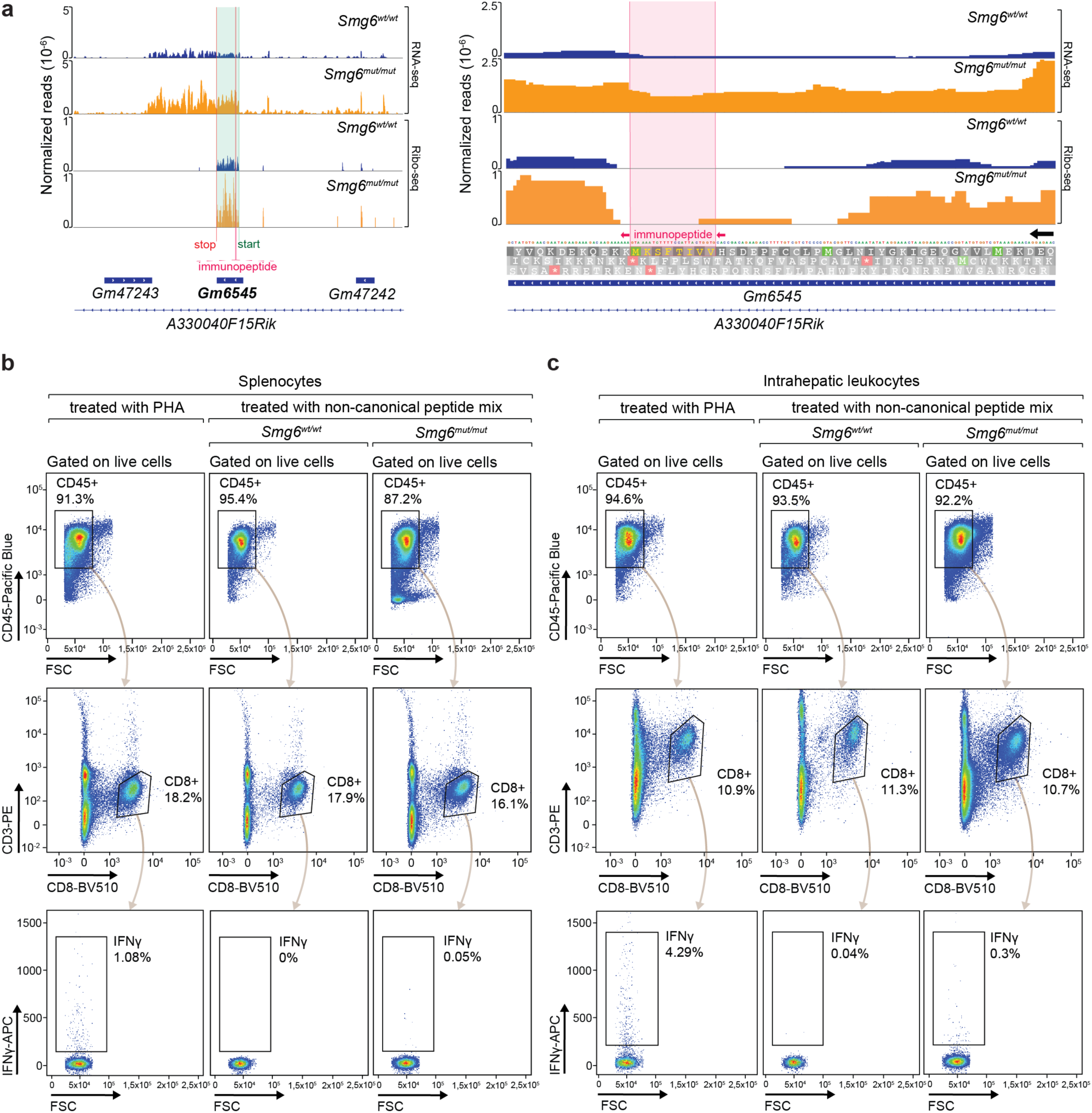
Increased non-canonical antigen expression in *Smg6* mutant livers. (a) RNA-seq (top panels) and Ribo-seq (bottom panels) read coverage plots show upregulated transcript and ribosome footprint abundance for *Gm6545* in *Smg6^mut/mut^*compared to *Smg6^wt/wt^*. Left panel shows the whole genomic region, including the CDS (highlighted in turquoise) where the non-canonical polypeptide is translated, while right panel zooms into the specific region where the immunopeptide is translated (highlighted in light magenta). Read coverage plots were generated by merging sequencing data from three biological replicates (individual livers) per group. (b, c) Representative flow cytometry gating strategies for one liver and one spleen from each group for IFNγ^+^CD8^+^ T-cells, treated with PHA (positive control) or non-canonical peptide pool.

**Supplementary Figure 5.**
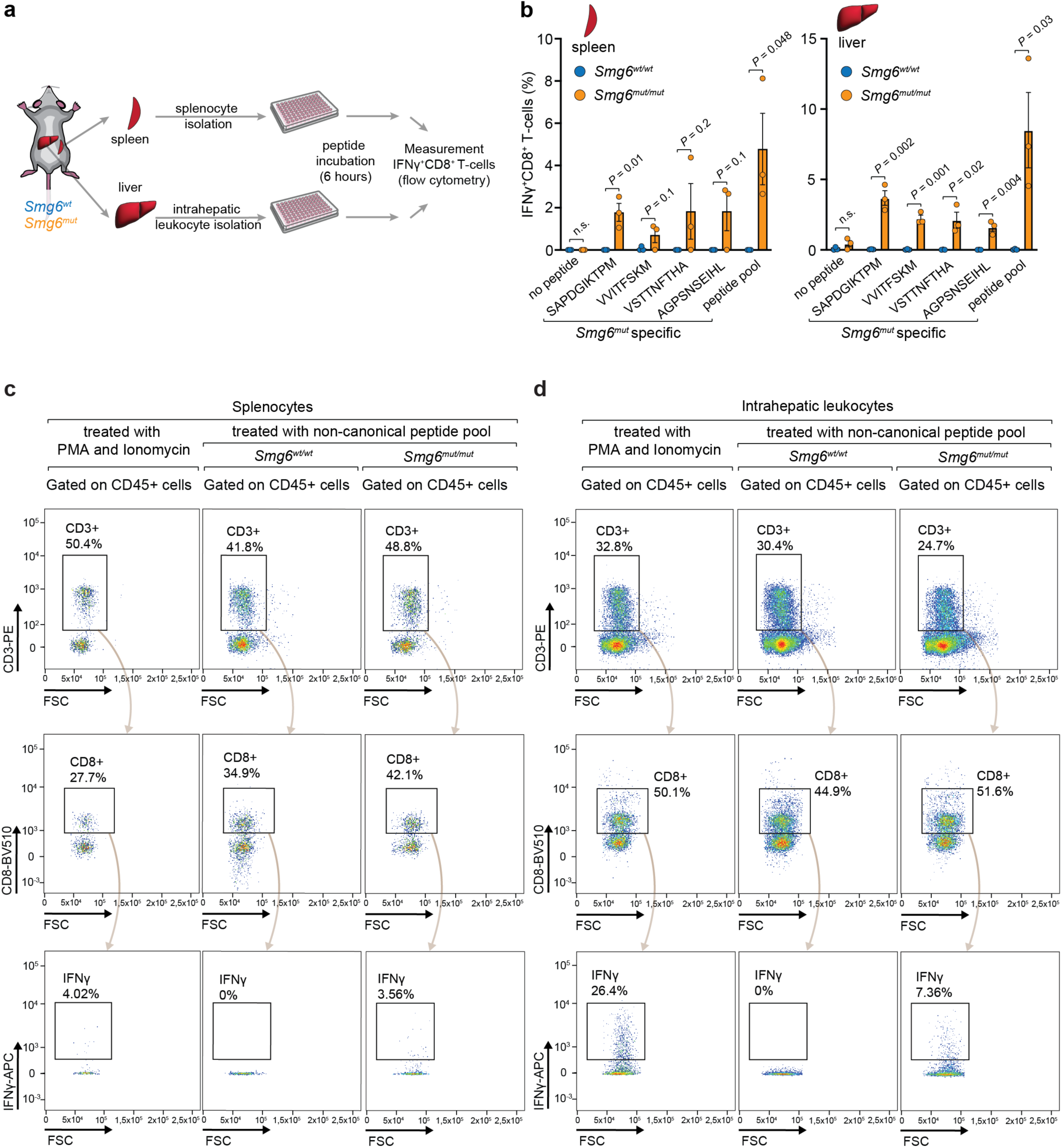
Testing the immunogenicity of *Smg6^mut^*-specific noncanonical antigens individually. (a) Schematic of the immunogenicity quantification experiment. *Smg6^wt/wt^* and *Smg6^mut/mut^* mice were sacrificed 5 weeks post-tamoxifen. Spleens and livers were extracted from the animals and leukocytes were isolated. The cells were then incubated with the *Smg6^mut^*-specific non-canonical peptides (SAPDGIKTPM, VVITFSKM, VSTTNFTHA or AGPSNSEIHL) separately or with a pool containing all these four peptides, for 6 hours. The percentages of IFNγ^+^CD8^+^ T-cells were used as a readout for the immunogenicity of each peptide pool (n = 3 mice for each group). (b) Overall IFNγ^+^CD8^+^ T-cell percentage per tissue (left panel, spleen; right panel, liver), as assessed by flow cytometry, upon treatment with the indicated *Smg6^mut^*-specific non-canonical peptides separately of with a pool containing all these four peptides. Bars indicate means, and the error bars SD; individual datapoints are measurements from independent animals (n = 3 mice for each group). The *P*-values were calculated with two-tailed unpaired *t*-test. (c, d) Representative flow cytometry gating strategies for one liver and one spleen from each group for IFNγ^+^CD8^+^ T-cells, treated with PMA and Ionomycin (positive control) or non-canonical peptide pool.

**Supplementary Figure 6.**
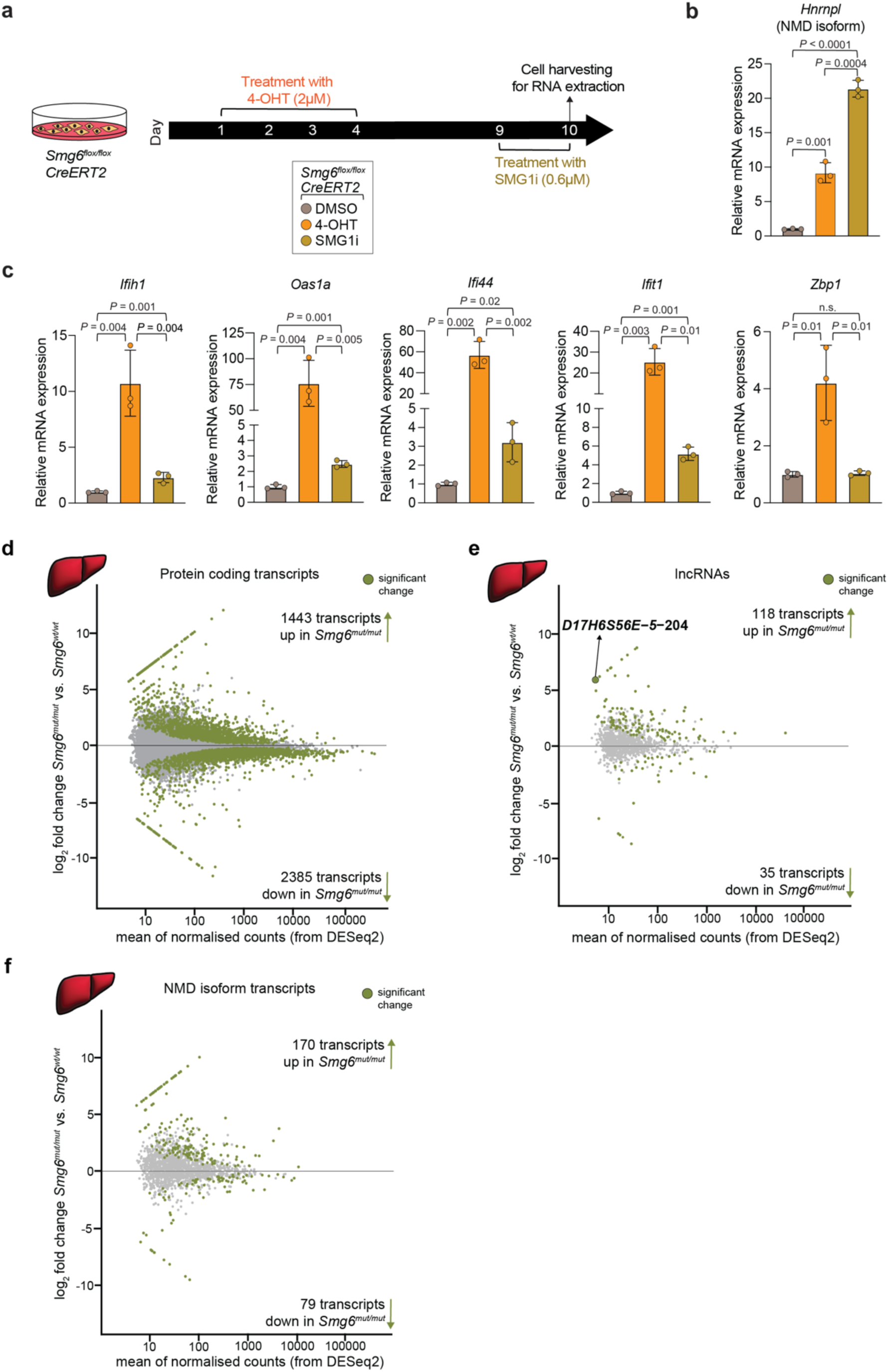
Comparison of the ISG expression via *Smg6* mutation vs. SMG1 inhibition. (a) Schematic of the experiment to analyze ISG expression in primary mouse fibroblasts. To induce *Smg6* mutation in *Smg6^flox/flox^*; *CreERT2* fibroblasts, cells were treated with 2 µM 4-hydroxytamoxifen (4-OHT) during days 1 to 4, to inhibit SMG1 cells were treated with 0.6 µM SMG1 inhibitor (SMG1i) at day 9. All cells were collected at day 10 for RNA isolation and qPCR. Control treatment was DMSO. (b) Relative mRNA expression of the endogenous, diagnostic NMD-target, an *Hnrnpl* splice form, in samples collected as described in (a). Three biological replicates were used per group. (c) Relative mRNA expression of indicated ISGs in same samples as in (b). Three biological replicates were used per group. (d-f) Differential expression analysis of liver transcriptomics data for protein coding transcripts (d), lncRNAs (e), and annotated NMD isoform transcripts (f). Significantly changed transcripts are highlighted in olive. For (b) and (c) data are plotted as means, and the error bars indicate SD. The *P*-values were calculated with two-tailed unpaired *t*-test.

**Supplementary Figure 7.**
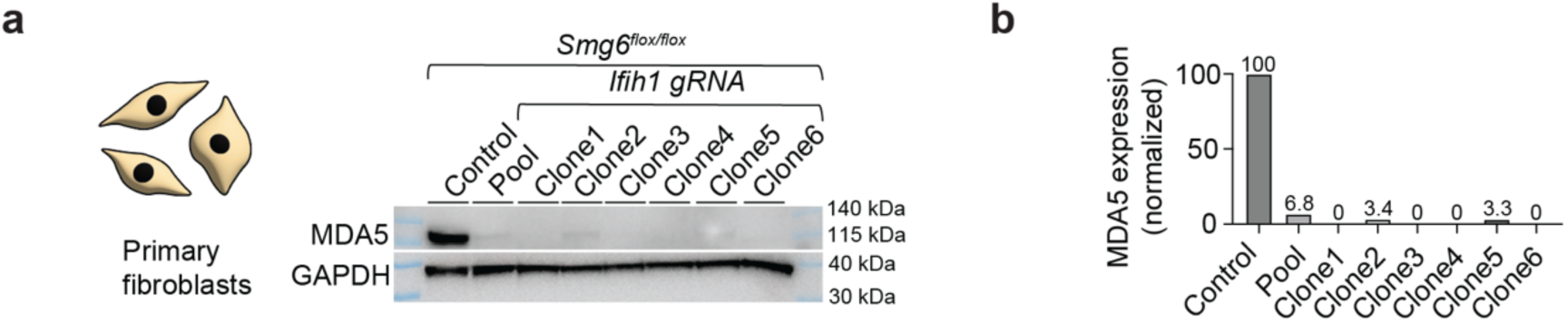
Deletion of *Ifih1* in *Smg6^flox/flox^* fibroblasts via CRISPR. (a) Western blot confirmation of the loss of MDA5 activity in different fibroblast single-cell clones. GAPDH serves as loading control. (b) Quantification of western blot from (a).

**Supplementary Figure 8.**
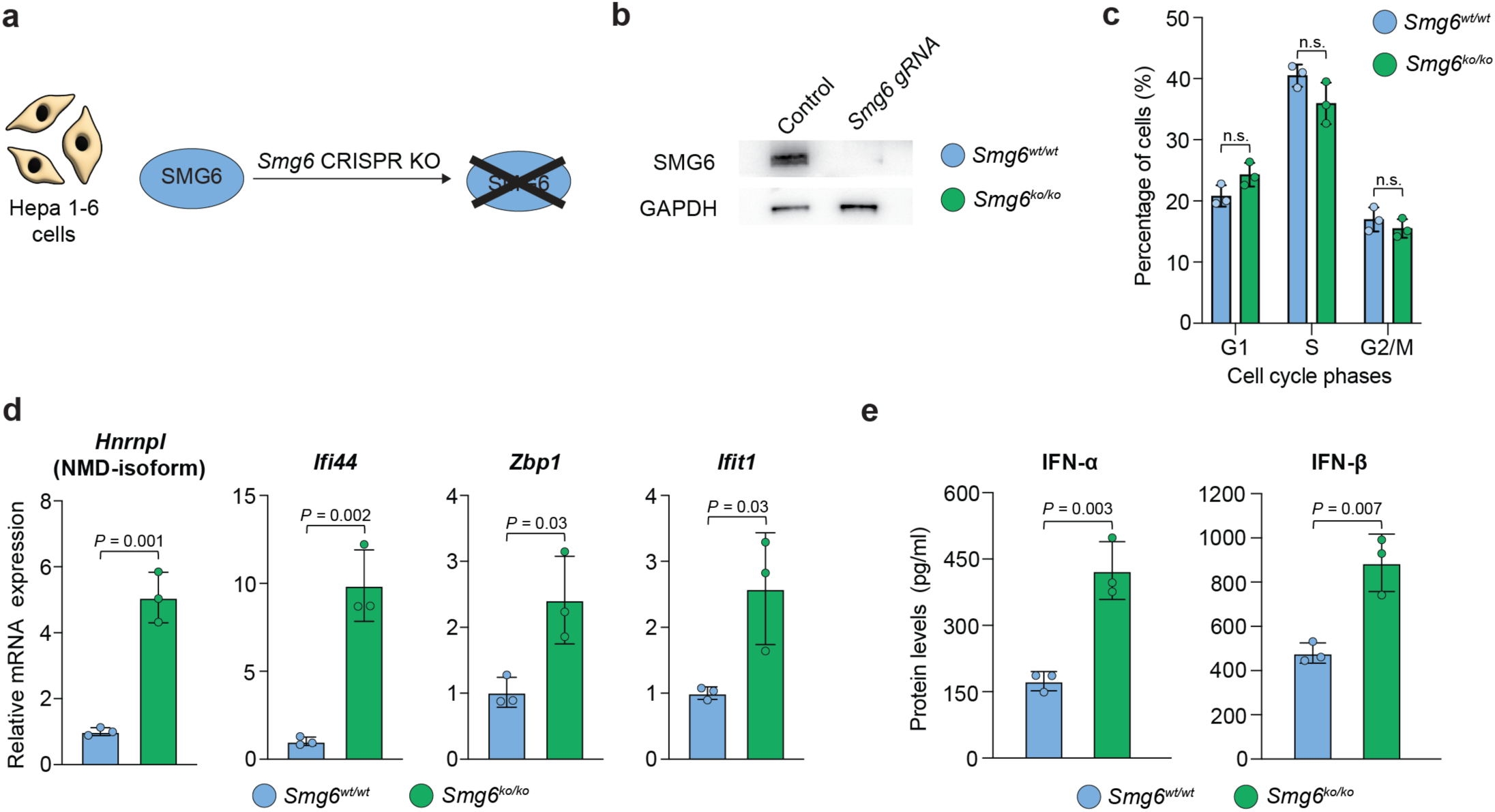
*Smg6^ko^* leads to activation of type I IFN response pathway in Hepa 1-6 cells. (a) Schematic of the *Smg6* knockout in Hepa1-6 cells. (b) Western blot validation of the absence of SMG6 expression. GAPDH served as loading control. (c) Analysis of cell cycle distribution via flow cytometry using propidium iodide staining. Three biological replicates were used per group. (d) Relative mRNA expression of the endogenous, diagnostic NMD-target, an *Hnrnpl* splice form and indicated ISGs. Three biological replicates were used per group. (e) ELISA analysis of IFN-α and IFN-β. Hepa1-6 cells (*Smg6^wt/wt^* or *Smg6^ko/ko^*) were induced with 1 µM of poly I:C for 8 hours. The supernatants were collected 48 hours post-treatment, and protein levels were detected via ELISA. Three biological replicates were used per group. For (c), (d), and (e) data are plotted as means, and the error bars indicate SD. The *P*-values were calculated with two-tailed unpaired *t*-test.

**Supplementary Figure 9.**
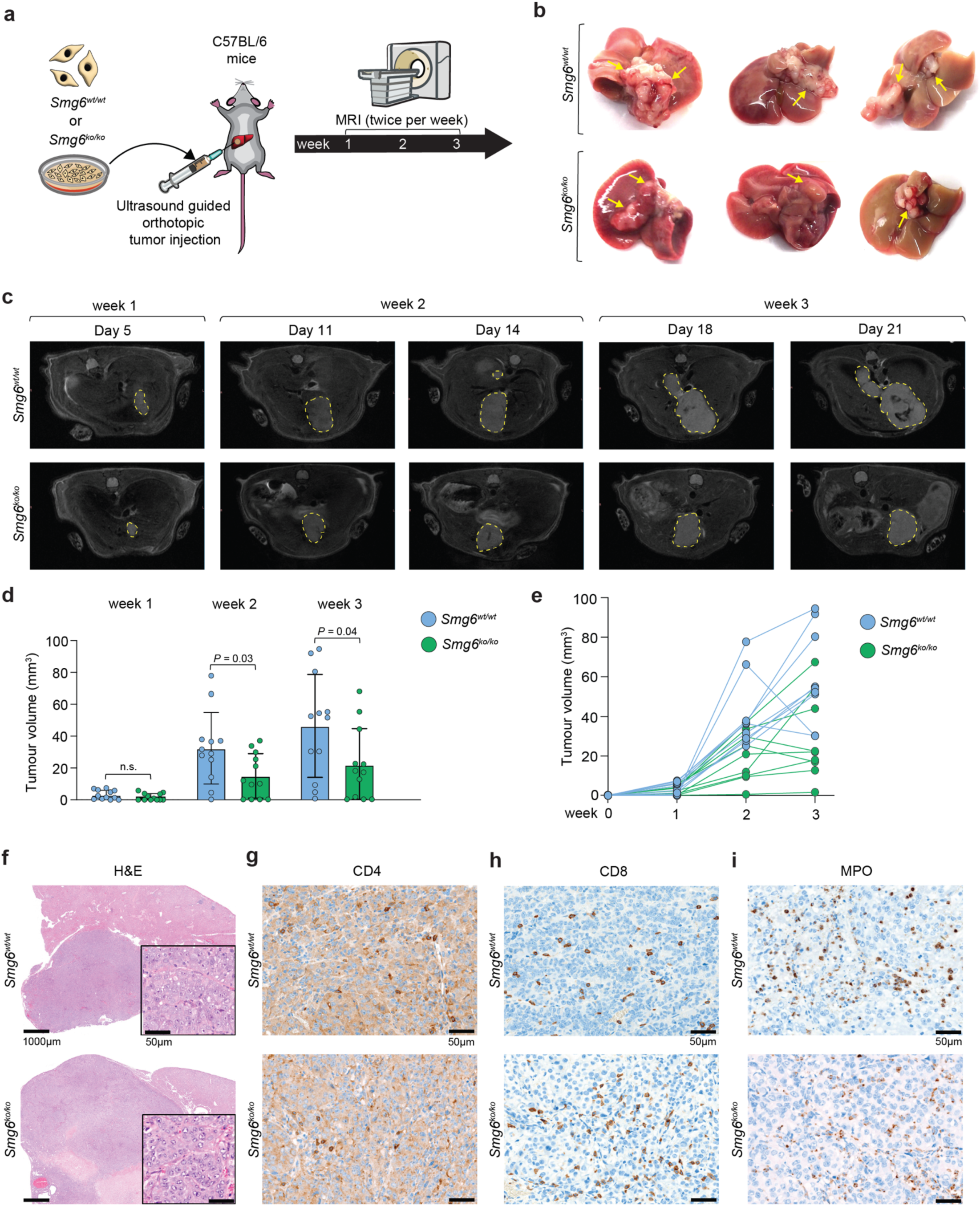
Tumours arising from orthotopically implanted *Smg6^ko/ko^* cells exhibit reduced growth compared with *Smg6^wt/wt^* counterparts. (a) Schematic of the ultrasound-guided orthotopic Hepa1-6 tumour injection (*Smg6^wt/wt^* and *Smg6^ko/ko^*) to the immunocompetent C57BL/6 mice. Tumour progression was then evaluated via MRI twice per week until the end of week 3 (n = 12 mice per group). (b) Representative whole liver images of three livers from each group. Tumours are indicated with yellow arrows. (c) Representative longitudinal abdominal MRI scans for one representative mouse each, orthotopically transplanted with *Smg6^wt/wt^* or *Smg6^ko/ko^* Hepa1-6 cells. Tumours in liver are marked by yellow dashed lines. Data are plotted as means, and the error bars indicate SD. The *P*-values were calculated with two-tailed unpaired *t*-test. (d) Longitudinal tumour volume measurement from 30 slices for each MRI (n = 12 mice for each group). (e) Trajectory of tumour volume evolution of 9 mice from each group that showed the highest tumour volume. (f) Representative H&E images from *Smg6^wt/wt^* and *Smg6^ko/ko^* liver tumours at magnifications 2x (main images) and 40x (inserts). Similar tumour morphology is observed between two genotypes. The tumour cells in both groups are large and polymorphic, with abundant basophilic cytoplasm, a large round nucleus, open chromatin, and a prominent central nucleolus. (g) Representative IHC images for CD4 staining from *Smg6^wt/wt^* and *Smg6^ko/ko^* liver tumours at 40x magnification. These images are part of a larger IHC analysis that was performed on a randomly chosen subset of liver samples from the cohort that showed clear liver engraftment of the tumours (*Smg6^wt/wt^*, n = 4; *Smg6^ko/ko^*, n = 3). For each of the livers, the number of CD4^+^ T-cells was counted in 10 randomly selected fields. Quantitative comparison revealed that CD4^+^ T-cell infiltration was higher in *Smg6^ko/ko^* tumours (median 14.25 cells/field) than in tumours of *Smg6^wt/wt^* origin (median 5.25 cells/field). However, due to the relatively low sample number and variabilities across individuals, statistical significance was not reached. Of note, in particular one sample from the *Smg6^wt/wt^* group exhibited exceptionally high numbers of both CD4^+^ and CD8^+^ T-cells even in normal parenchymal regions, indicating a possible lymphocyte infiltration issue caused by other reasons that then tumour; although likely an outlier, we did not exclude this individual from our overall analysis. (h) As in panel (g), representative IHC images for CD8. CD8^+^ T-cell infiltration was higher in tumours of *Smg6^ko/ko^* origin (median 74.5 cells/field) than in those of *Smg6^wt/wt^* origin (median 47.5 cells/field); despite this large difference, as explained for panel (g), due to the relatively low sample number and variabilities across individuals, statistical significance was not reached. (i) As in (g) and (h), representative IHC images for MPO from both *Smg6^wt/wt^* and *Smg6^ko/ko^* liver tumours at 40x magnification. Our larger analysis (*Smg6^wt/wt^*, n = 4; *Smg6^ko/ko^*, n = 3) revealed that the infiltration by granulocytes in the normal liver parenchyma was low in both groups. Within tumour regions, increased MPO-positive infiltration was observed in both genotypes, yet the calculated median scores were near-identical between *Smg6^wt/wt^* and *Smg6^ko/ko^* (scores 2.25 vs. 2).

**Supplementary Figure 10.**
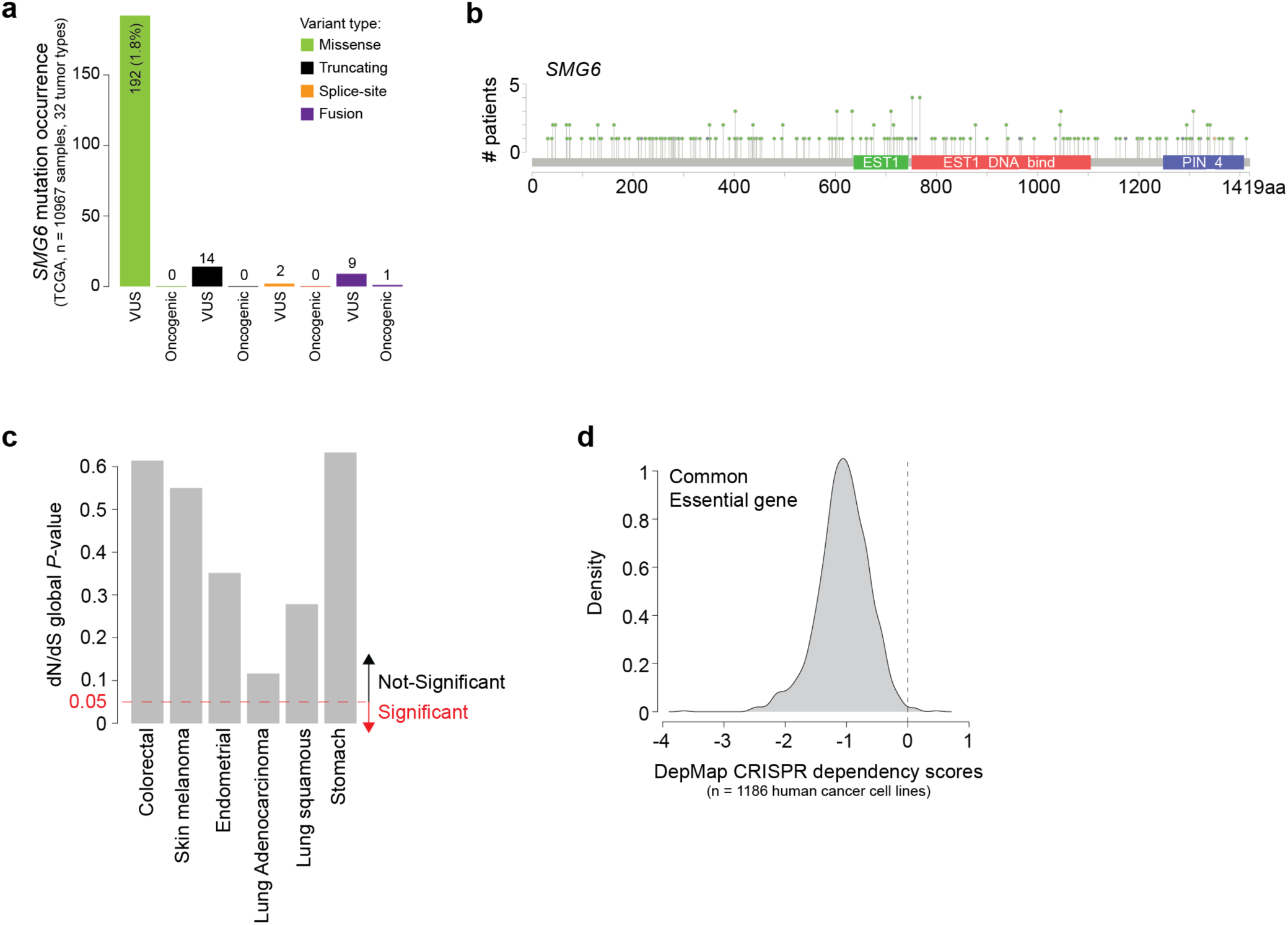
Exploring *SMG6* mutations in human cancer using TCGA data. a) The barplot reports the number of mutations observed in the TCGA Pan-cancer atlas cohort ^59^, split by variant type and variant classification (VUS, variant of unknown significance; Oncogenic, predicted or demonstrated to be a cancer driver). Data and classifications are taken from the <u>cBioPortal</u> for cancer genomics (n = 10967 samples, 32 different tumour types). From this analysis, it can be concluded that mutations are overall very rare (< 2%), with the majority being passenger missense mutations. b) The lollipop plot shows the position in the protein sequence of the *SMG6* mutations in the TCGA dataset. Data and classifications are taken from the <u>cBioPortal</u> for cancer genomics. From this analysis, it can be concluded that missense mutations are rather uniformly scattered along the protein sequence, further supporting them being functionally neutral (i.e., functional/oncogenic missense mutations typically cluster within specific domains or are over-represented at a specific “hotspot” residue; lack of such patterns is taken as lack of positive selection). c) Analysis of non-synonymous and synonymous *SMG6* mutations via dN/dS ^61^ ratio using the tumour types where at least 10 mutated samples were found for *SMG6*. Barplot shows the *P*-values for *SMG6* in each tumour type. We conclude that *SMG6* is never found significantly recurrently mutated, further supporting lack of positive selection. d) <u>DepMap</u> dependency scores for *SMG6*. Briefly, DepMap consists of large-scale CRISPR screenings in human cancer cell lines (n = 1186 human cancer cell lines). Dependency scores quantify changes of viability/fitness measured in each cell line upon KO of a given gene. Negative values (in particular below −0.5) indicate loss of viability. The density plot shows the distribution of scores for SMG6. This distribution pattern indicates that SMG6 loss is deleterious across most cancer cell lines.

## References

1. Kurosaki, T., Popp, M.W. & Maquat, L.E. Quality and quantity control of gene expression by nonsense-mediated mRNA decay. Nature reviews. Molecular cell biology 20, 406–420 (2019).

2. Karousis, E.D. & Muhlemann, O. The broader sense of nonsense. Trends in biochemical sciences 47, 921–935 (2022).

3. Boehm, V. et al. SMG5-SMG7 authorize nonsense-mediated mRNA decay by enabling SMG6 endonucleolytic activity. Nature communications 12, 3965 (2021).

4. Arpa, E.S., Taschner, M., De Matos, M., Jonas, S. & Gatfield, D. Active Site Assembly by SMG5 as a Mechanism for SMG6 Endonuclease Licencing in Nonsense-mediated mRNA Decay. J Mol Biol, 169734 (2026).

5. Kurscheidt, K. et al. Composite SMG5-SMG6 PIN domain formation is essential for NMD. Nature communications 17 (2026).

6. Modena, M.S. et al. The PIN domain of SMG-5 functionally interacts with SMG-6 to stimulate NMD. Rna (2026).

7. Supek, F., Lehner, B. & Lindeboom, R.G. To NMD or Not To NMD: nonsense-mediated mRNA decay in cancer and other genetic diseases. Trends in Genetics 37, 657–668 (2021).

8. Lindeboom, R.G.H., Vermeulen, M., Lehner, B. & Supek, F. The impact of nonsense-mediated mRNA decay on genetic disease, gene editing and cancer immunotherapy. Nat Genet 51, 1645–1651 (2019).

9. Wu, C.C. et al. Immuno-genomic landscape of osteosarcoma. Nature communications 11, 1008 (2020).

10. Litchfield, K. et al. Escape from nonsense-mediated decay associates with anti-tumor immunogenicity. Nature communications 11, 3800 (2020).

11. Oka, M. et al. Aberrant splicing isoforms detected by full-length transcriptome sequencing as transcripts of potential neoantigens in non-small cell lung cancer. Genome biology 22, 9 (2021).

12. Becker, J.P. et al. NMD inhibition by 5-azacytidine augments presentation of immunogenic frameshift-derived neoepitopes. iScience 24, 102389 (2021).

13. Zhang, Y. et al. Immunotherapy for breast cancer using EpCAM aptamer tumor-targeted gene knockdown. Proceedings of the National Academy of Sciences of the United States of America 118 (2021).

14. Pastor, F., Kolonias, D., Giangrande, P.H. & Gilboa, E. Induction of tumour immunity by targeted inhibition of nonsense-mediated mRNA decay. Nature 465, 227–230 (2010).

15. Meraviglia-Crivelli, D. et al. IL-6/STAT3 signaling in tumor cells restricts the expression of frameshift-derived neoantigens by SMG1 induction. Mol Cancer 21, 211 (2022).

16. Meraviglia-Crivelli, D. et al. A pan-tumor-siRNA aptamer chimera to block nonsense-mediated mRNA decay inflames and suppresses tumor progression. Mol Ther Nucleic Acids 29, 413–425 (2022).

17. Nasif, S., et al. Inhibition of nonsense-mediated mRNA decay reduces the tumorigenicity of human fibrosarcoma cells. NAR Cancer 5, zcad048 (2023).

18. Hwang, J. & Maquat, L.E. Nonsense-mediated mRNA decay (NMD) in animal embryogenesis: to die or not to die, that is the question. Curr Opin Genet Dev 21, 422–430 (2011).

19. Li, T. et al. Smg6/Est1 licenses embryonic stem cell differentiation via nonsense-mediated mRNA decay. The EMBO journal 34, 1630–1647 (2015).

20. Katsioudi, G. et al. A conditional Smg6 mutant mouse model reveals circadian clock regulation through the nonsense-mediated mRNA decay pathway. Sci Adv 9, eade2828 (2023).

21. Guri, Y. et al. mTORC2 Promotes Tumorigenesis via Lipid Synthesis. Cancer Cell 32, 807–823 e812 (2017).

22. Mossmann, D. et al. Arginine reprograms metabolism in liver cancer via RBM39. Cell 186, 5068–5083 e5023 (2023).

23. Postic, C. & Girard, J. The role of the lipogenic pathway in the development of hepatic steatosis. Diabetes Metab 34, 643–648 (2008).

24. Snoek, M., van Dinten, L. & van Vugt, H. A novel gene, G7e, resembling a viral envelope gene, is located at the recombinational hot spot in the class III region of the mouse MHC. Genomics 38, 5–12 (1996).

25. Schoggins, J.W. et al. A diverse range of gene products are effectors of the type I interferon antiviral response. Nature 472, 481–485 (2011).

26. Gopalsamy, A. et al. Identification of pyrimidine derivatives as hSMG-1 inhibitors. Bioorg Med Chem Lett 22, 6636–6641 (2012).

27. Sung, J.H. et al. Chemokine guidance of central memory T cells is critical for antiviral recall responses in lymph nodes. Cell 150, 1249–1263 (2012).

28. Bowling, E.A. et al. Spliceosome-targeted therapies trigger an antiviral immune response in triple-negative breast cancer. Cell 184, 384–403 e321 (2021).

29. Zheng, R. et al. hnRNPM protects against the dsRNA-mediated interferon response by repressing LINE-associated cryptic splicing. Mol Cell 84, 2087–2103 e2088 (2024).

30. Dias Junior, A.G., Sampaio, N.G. & Rehwinkel, J. A Balancing Act: MDA5 in Antiviral Immunity and Autoinflammation. Trends Microbiol 27, 75–85 (2019).

31. Lewis, B.P., Green, R.E. & Brenner, S.E. Evidence for the widespread coupling of alternative splicing and nonsense-mediated mRNA decay in humans. Proceedings of the National Academy of Sciences 100, 189–192 (2003).

32. Zhou, H. & Deng, X.W. Intron Retention, an Orchestrated Program of Gene Expression Regulation. Bioessays 47, e202400248 (2025).

33. Rehwinkel, J. & Gack, M.U. RIG-I-like receptors: their regulation and roles in RNA sensing. Nat Rev Immunol 20, 537–551 (2020).

34. Pichlmair, A. et al. Activation of MDA5 requires higher-order RNA structures generated during virus infection. J Virol 83, 10761–10769 (2009).

35. Schonborn, J. et al. Monoclonal antibodies to double-stranded RNA as probes of RNA structure in crude nucleic acid extracts. Nucleic Acids Res 19, 2993–3000 (1991).

36. Pepin, G. et al. Cre-dependent DNA recombination activates a STING-dependent innate immune response. Nucleic Acids Res 44, 5356–5364 (2016).

37. Darlington, G.J., Bernhard, H.P., Miller, R.A. & Ruddle, F.H. Expression of liver phenotypes in cultured mouse hepatoma cells. J Natl Cancer Inst 64, 809–819 (1980).

38. Laky, K. & Kruisbeek, A.M. In Vivo Depletion of T Lymphocytes. Curr Protoc Immunol 113, 4 1 1–4 1 9 (2016).

39. Holicek, P. et al. Type I interferon and cancer. Immunol Rev 321, 115–127 (2024).

40. Rehwinkel, J. & Mehdipour, P. ADAR1: from basic mechanisms to inhibitors. Trends Cell Biol 35, 59–73 (2025).

41. Rigby, R.E. & Rehwinkel, J. RNA degradation in antiviral immunity and autoimmunity. Trends Immunol 36, 179–188 (2015).

42. Johnson, J.L. et al. Inhibition of Upf2-Dependent Nonsense-Mediated Decay Leads to Behavioral and Neurophysiological Abnormalities by Activating the Immune Response. Neuron 104, 665–679 e668 (2019).

43. Schuler, M., Dierich, A., Chambon, P. & Metzger, D. Efficient temporally controlled targeted somatic mutagenesis in hepatocytes of the mouse. Genesis 39, 167–172 (2004).

44. Horie, Y. et al. Hepatocyte-specific Pten deficiency results in steatohepatitis and hepatocellular carcinomas. J Clin Invest 113, 1774–1783 (2004).

45. Kwiatkowski, D.J. et al. A mouse model of TSC1 reveals sex-dependent lethality from liver hemangiomas, and up-regulation of p70S6 kinase activity in Tsc1 null cells. Hum Mol Genet 11, 525–534 (2002).

46. Benechet, A.P. et al. Dynamics and genomic landscape of CD8(+) T cells undergoing hepatic priming. Nature 574, 200–205 (2019).

47. Gatfield, D. et al. Integration of microRNA miR-122 in hepatic circadian gene expression. Genes Dev 23, 1313–1326 (2009).

48. Dobin, A. et al. STAR: ultrafast universal RNA-seq aligner. Bioinformatics 29, 15–21 (2013).

49. Love, M.I., Huber, W. & Anders, S. Moderated estimation of fold change and dispersion for RNA-seq data with DESeq2. Genome Biol 15, 550 (2014).

50. Snieckute, G. et al. ROS-induced ribosome impairment underlies ZAKalpha-mediated metabolic decline in obesity and aging. Science 382, eadf3208 (2023).

51. Martin, M. Cutadapt removes adapter sequences from high-throughput sequencing reads. EMBnet.journal 17, 10–12 (2011).

52. Lawrence, M. et al. Software for computing and annotating genomic ranges. PLoS Comput Biol 9, e1003118 (2013).

53. Chong, C. et al. High-throughput and Sensitive Immunopeptidomics Platform Reveals Profound Interferongamma-Mediated Remodeling of the Human Leukocyte Antigen (HLA) Ligandome. Mol Cell Proteomics 17, 533–548 (2018).

54. Kong, A.T., Leprevost, F.V., Avtonomov, D.M., Mellacheruvu, D. & Nesvizhskii, A.I. MSFragger: ultrafast and comprehensive peptide identification in mass spectrometry-based proteomics. Nat Methods 14, 513–520 (2017).

55. Gfeller, D. et al. Improved predictions of antigen presentation and TCR recognition with MixMHCpred2.2 and PRIME2.0 reveal potent SARS-CoV-2 CD8(+) T-cell epitopes. Cell Syst 14, 72–83 e75 (2023).

56. Unlu, B. et al. Global analysis of suppressor mutations that rescue human genetic defects. Genome Med 15, 78 (2023).

57. de Faria, I.J.S., Imler, J.L. & Marques, J.T. Protocol for the analysis of double-stranded RNAs in virus-infected insect cells using anti-dsRNA antibodies. STAR Protoc 4, 102033 (2023).

58. Smit, A., Hubley, R. & Green, P. RepeatMasker Open-3.0. http://www.repeatmasker.org (1996-2004).

59. Sanchez-Vega, F. et al. Oncogenic Signaling Pathways in The Cancer Genome Atlas. *Cell* 173, 321-337 e310 (2018).

60. Cerami, E. et al. The cBio cancer genomics portal: an open platform for exploring multidimensional cancer genomics data. Cancer Discov 2, 401–404 (2012).

61. Martincorena, I. et al. Universal Patterns of Selection in Cancer and Somatic Tissues. Cell 171, 1029–1041 e1021 (2017).

62. Arafeh, R., Shibue, T., Dempster, J.M., Hahn, W.C. & Vazquez, F. The present and future of the Cancer Dependency Map. Nat Rev Cancer 25, 59–73 (2025).

63. Edgar, R., Domrachev, M. & Lash, A.E. Gene Expression Omnibus: NCBI gene expression and hybridization array data repository. Nucleic Acids Res 30, 207–210 (2002).

64. Perez-Riverol, Y. et al. The PRIDE database resources in 2022: a hub for mass spectrometry-based proteomics evidences. Nucleic Acids Res 50, D543–D552 (2022).

